# RIT2 reduces LRRK2 kinase activity and protects against alpha-synuclein neuropathology

**DOI:** 10.1101/2020.10.21.348144

**Authors:** Julia Obergasteiger, Anne-Marie Castonguay, Giulia Frapporti, Evy Lobbestael, Veerle Baekelandt, Andrew A. Hicks, Peter P. Pramstaller, Claude Gravel, Corrado Corti, Martin Lévesque, Mattia Volta

## Abstract

In Parkinson’s disease (PD) misfolded alpha-synuclein (aSyn) accumulates in the substantia nigra, where dopaminergic neurons are progressively lost. The mechanisms underlying aSyn pathology are still unclear but hypothesized to involve the autophagy-lysosome pathway (ALP). LRRK2 mutations are a major cause of familial and sporadic PD, hyperactivate kinase activity and its pharmacological inhibition reduces pS129-aSyn inclusions. We observed selective downregulation of the novel PD risk factor *RIT2* in G2019S-LRRK2 expressing cells. Here we studied whether RIT2 could modulate LRRK2 kinase activity. RIT2 overexpression in G2019S-LRRK2 cells rescued ALP abnormalities and diminished aSyn inclusions. *In vivo*, viral mediated overexpression of RIT2 conferred neuroprotection against AAV-A53T-aSyn. Furthermore, RIT2 overexpression prevented the A53T-aSyn-dependent increase of LRRK2 kinase activity *in vivo*. Our data indicate that RIT2 inhibits overactive LRRK2 to ameliorate ALP impairment and counteract aSyn aggregation and related deficits. Targeting RIT2 could represent a novel strategy to combat neuropathology in familial and idiopathic PD.

## Introduction

Parkinson’s disease (PD) is the second most common neurodegenerative disorder affecting 2-3% of people over the age of 65. The disease is characterized by motor symptoms including resting tremor, rigidity, bradykinesia, and postural instability, which originate from the loss of dopaminergic (DA) neurons in the substantia nigra pars compacta (SNc) (Poewe et al., 2017; Tysnes & Storstein, 2017). Around 15% of PD patients have a family history and 5-10% of cases are caused by mutations and alterations in specific genes (e.g. SNCA, LRRK2, Parkin, VPS35) (Lesage & Brice, 2009). The vast majority of cases are sporadic and are thought to be caused by a combination of genetic and environmental factors (Dauer & Przedborski, 2003; Deng et al., 2018). Gene variants in alpha-synuclein (aSyn) and leucine-rich repeat kinase 2 (LRRK2) lead to autosomal dominant PD, mostly presenting intracytoplasmic protein aggregates called Lewy bodies (LBs). LBs are one of the main neuropathological hallmarks of PD and a group of other neurological disorders termed synucleinopathies, and are mainly composed of oligomeric and fibrillar aSyn (Dickson, 2018; Goedert et al., 2013). Mechanisms underlying the formation and clearance of these inclusions are still under intense investigation. The autophagy-lysosome pathway (ALP) is a conserved and tightly regulated cellular process leading to lysosomal degradation of long-lived proteins, defective organelles and protein aggregates. Cargo delivery to the lysosome can be achieved by three main pathways: microautophagy, chaperone-mediated autophagy (CMA) and macroautophagy. The latter involves the formation of a double-membrane vesicle, termed autophagosome, which engulfs the cargo. The autophagosome then fuses with the lysosome where final degradation takes place (Finkbeiner, 2020; Kenney & Benarroch, 2015). The ALP is dysregulated in PD and, amongst other cellular processes, likely contributes to the defective clearance of aSyn (Anglade et al., 1997; Dehay et al., 2010; Nixon, 2013).

LRRK2 is a multifunctional protein composed of a catalytic GTPase and kinase core and several protein binding domains (Paisan-Ruiz et al., 2013). LRRK2 is expressed in different tissues with high expression levels in the lung, kidney and white blood cells (Fuji et al., 2015). In the brain, LRRK2 is expressed ubiquitously and not restricted to neurons (Miklossy et al., 2006). Mutations in the catalytic core have been associated with late-onset familial PD, with the G2019S substitution leading to a 3-4-fold increase in kinase activity and being the most common genetic cause of PD (Trinh & Farrer, 2013). Cellular roles of LRRK2 are varied and include synaptic transmission, vesicle trafficking and autophagy (Beccano-Kelly et al., 2014; Beccano-Kelly et al., 2015; Parisiadou et al., 2014; Piccoli et al., 2011). Numerous studies have reported that mutant LRRK2 alters the ALP, but a common consensus on the specific effects and the impact of LRRK2 mutations on the ALP is currently lacking. (Manzoni & Lewis, 2017; Manzoni et al., 2013; Saez-Atienzar et al., 2014; Schapansky et al., 2018). Moreover, LRRK2 kinase inhibition is beneficial against aSyn neuropathology, suggesting that LRRK2 activity mediates accumulation of pathologic aSyn (Volpicelli-Daley et al., 2016).

Recent genome-wide association studies (GWAS) identified the locus containing the *RIT2* gene to be associated with increased risk for PD (Nalls et al., 2019; Nalls et al., 2014; Pankratz et al., 2012). *RIT2* encodes for the small GTPase Rin, which is mainly expressed in neural tissue (Lee et al., 1996). Rin belongs to the Ras superfamily of GTPases and is implicated in MAPK-mediated neurite outgrowth (Hoshino & Nakamura, 2003; Shi et al., 2005; Spencer et al., 2002), calcium signaling (Hoshino & Nakamura, 2003; Lee et al., 1996) and DA transporter (DAT) trafficking (Navaroli et al., 2011). Recently, we reviewed the possible regulation of autophagy by GTPase-MAPK pathways. Since *RIT2* modulates the activity of MAPKs (including p38 and ERK1/2), we hypothesized that it could contribute to the regulation of the ALP (Obergasteiger et al., 2018). Notably, LRRK2 also participates to MAPK signaling (Rui et al., 2018).

Here, we investigate the effects of enhanced *RIT2* expression on aSyn pathology, ALP regulation, and the underlying molecular mechanism using preclinical *in vitro* and *in vivo* PD models. Overexpression of *RIT2* in G2019S-LRRK2 neuroblastoma cells restored ALP deficits and reduced accumulation of phosphorylated aSyn (pS129-aSyn), phenocopying the effects of pharmacological LRRK2 kinase inhibition (Obergasteiger et al., 2020). The selective overexpression of *Rit2* in DA neurons in the mouse SNc attenuated nigrostriatal neurodegeneration and pS129-aSyn neuropathology induced by virally expressed aSyn. Importantly, we found that enhanced *RIT2* expression inhibits both recombinant (*in vitro*) and endogenous (*in vivo*) LRRK2 kinase activity.

## Results

### RIT2 gene expression is reduced in recombinant LRRK2 cells and in dopamine neurons from idiopathic PD patients

Genetic alterations in the *RIT2* promoter region have been associated to PD and are hypothesized to alter expression levels (Latourelle et al., 2012). Accordingly, *RIT2* expression was reduced in the SNc of PD patients (Bossers et al., 2009). Using publicly available datasets, we analyzed *RIT2* mRNA expression in human and rodent samples. The Geo dataset GSE20141 reports microarray data obtained from laser-capture microdissected SNc DA neurons from idiopathic PD patients and controls (Zheng et al., 2010). Consistent with literature, we found that *RIT2* expression is downregulated by about 2.2-fold in DA neurons from idiopathic PD patients, when compared to controls (Fig. 1A). Interestingly, *RIT2* mRNA levels in total brain tissue were not changed (GSE7621 (Lesnick et al., 2007); Fig. S1A), indicating a specific downregulation of *RIT2* in DA neurons of the SNc. *In vitro, RIT2* mRNA levels were also reduced in DA neurons generated from iPSCs carrying the A53T mutation in aSyn when compared to isogenic control cells (GSE46798 (Ryan et al., 2013); Fig. 1B) and in SK-N-SH neuroblastoma cells overexpressing A53T-aSyn (Fig. S1B) when compared to naïve cells.

**Figure 1:**
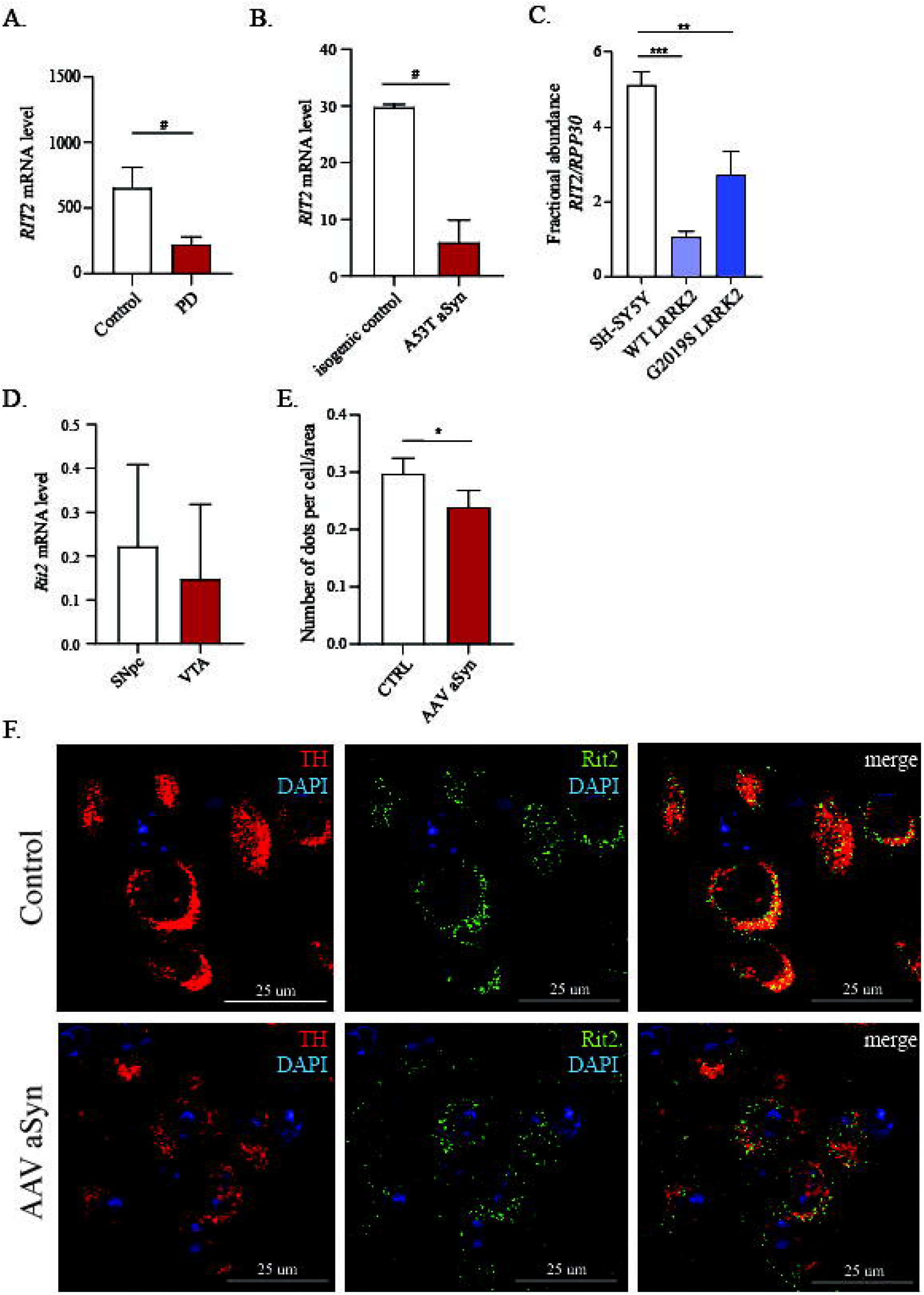
RIT2 is expressed in human and mouse brain, and is reduced in both sporadic PD patients and in cells overexpressing LRRK2. A) Geo dataset comparing *RIT2* expression levels in DA neurons of the SNc isolated by laser capture microdissection. mRNA levels were reduced by 2.2-fold in DA neurons from sporadic PD patients, when compared to healthy controls (GSE20141, controls=8, PD=10). # p<0.05, two-tailed Mann-Whitney test. B) Geo dataset (GSE46798) comparing *RIT2* expression in iPSC-derived DA neurons with A53T-aSyn mutation to control mutation-corrected neurons. # p<0.05 unpaired t-test with Welch’s correction. C) Droplet Digital PCR was carried out to assess *RIT2* mRNA levels in recombinant neuroblastoma cell lines. Fractional abundance of *RIT2* mRNA, normalized to RPP30, is reduced in WT and G2019S LRRK2 cells (n=3). Data are means±SEM of 3 independent experiments for ddPCR. **p<0.01, ***p<0.001, one-way ANOVA followed by Bonferroni’s post-hoc test. D) Geo dataset (GSE17542) comparing *Rit2* expression in SNc and VTA from TH-GFP mice. E-F) Fluorescent in situ hybridization (RNAscope^®^) was employed to show *Rit2* mRNA expression in DA neurons of the mouse SNc. E) AAV aSyn overexpression decreases Rit2 mRNA levels in mouse SNc. F) Neurons positive for *TH* mRNA display a robust level of *Rit2* mRNA in control animals, which is decreased with AAV aSyn overexpression.

To complement these analyses, we measured *RIT2* mRNA levels in WT-and G2019S-LRRK2 overexpressing SH-SY5Y neuroblastoma cells using droplet digital PCR (ddPCR) and found reduced gene expression in both cell lines, when compared to naïve SH-SY5Y cells (Fig. 1C). In rodents, *Rit2* was reported to be expressed in the brain (Zhou et al., 2011). The analysis of Geo dataset GSE17542 (Phani et al., 2010), comparing *Rit2* mRNA levels from laser-capture microdissected mouse SNc and ventral tegmental area (VTA) did not reveal significant regional differences in *Rit2* mRNA expression (Fig. 1D). To confirm *Rit2* expression in DA neurons of the mouse SNc, we performed multiplex *in situ* hybridization (RNAscope^®^ ISH technology) on mouse midbrain sections. High transcriptional levels of *Rit2* were found in DA (TH+) neurons of both the SNc and the VTA (Fig. 1F). Interestingly, AAV-A53T aSyn overexpression, inducing a loss of TH-positive neurons (see Fig. 5), significantly reduced *Rit2* mRNA levels (Fig 1E-F). Altogether, our results show that *RIT2* is expressed in both mouse and human DA neurons, and its expression is reduced in DA neurons from idiopathic PD patients as well as in cellular and mouse models of familial PD.

**Figure 2:**
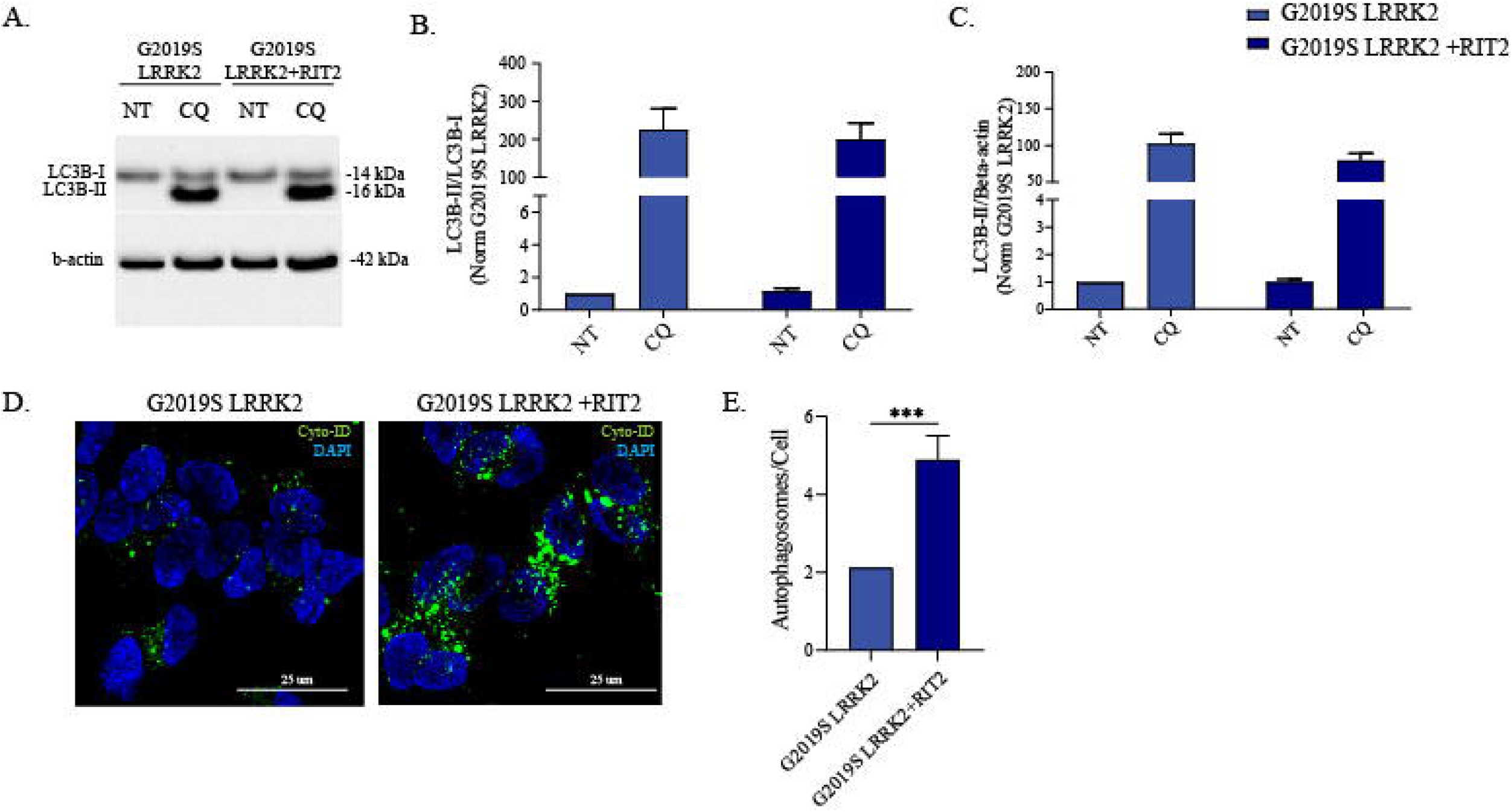
Overexpression of RIT2 does not affect autophagy initiation, but reduces autophagosome number. A) The autophagic flux was assessed in G2019S-LRRK2 and G2019S LRRK2+*RIT2* cells upon treatment with CQ (100 μM, 3h) and WB for LC3B. B) The ratio between LC3B-II and LC3B-I was not different, suggesting no differences in autophagic flux (n=3). C) Quantification of LC3B levels (normalized to β-actin) indicated no differences upon *RIT2* overexpression or CQ-treatment (n=3). D) Cyto-ID dye was used to visualize the distribution of autophagosomes. E) Quantification of Cyto-ID-positive vesicles revealed a significant increase in G2019S-LRRK2 cells, when *RIT2* was overexpressed (n=4). Data are means±SEM of 3 independent experiments for WB. In imaging experiments, 4 independent biological replicates were performed, and analysis conducted on 700-1000 cells per group in each experiment. ***p<0.001, unpaired two tailed Student’s t-test.

**Figure 3:**
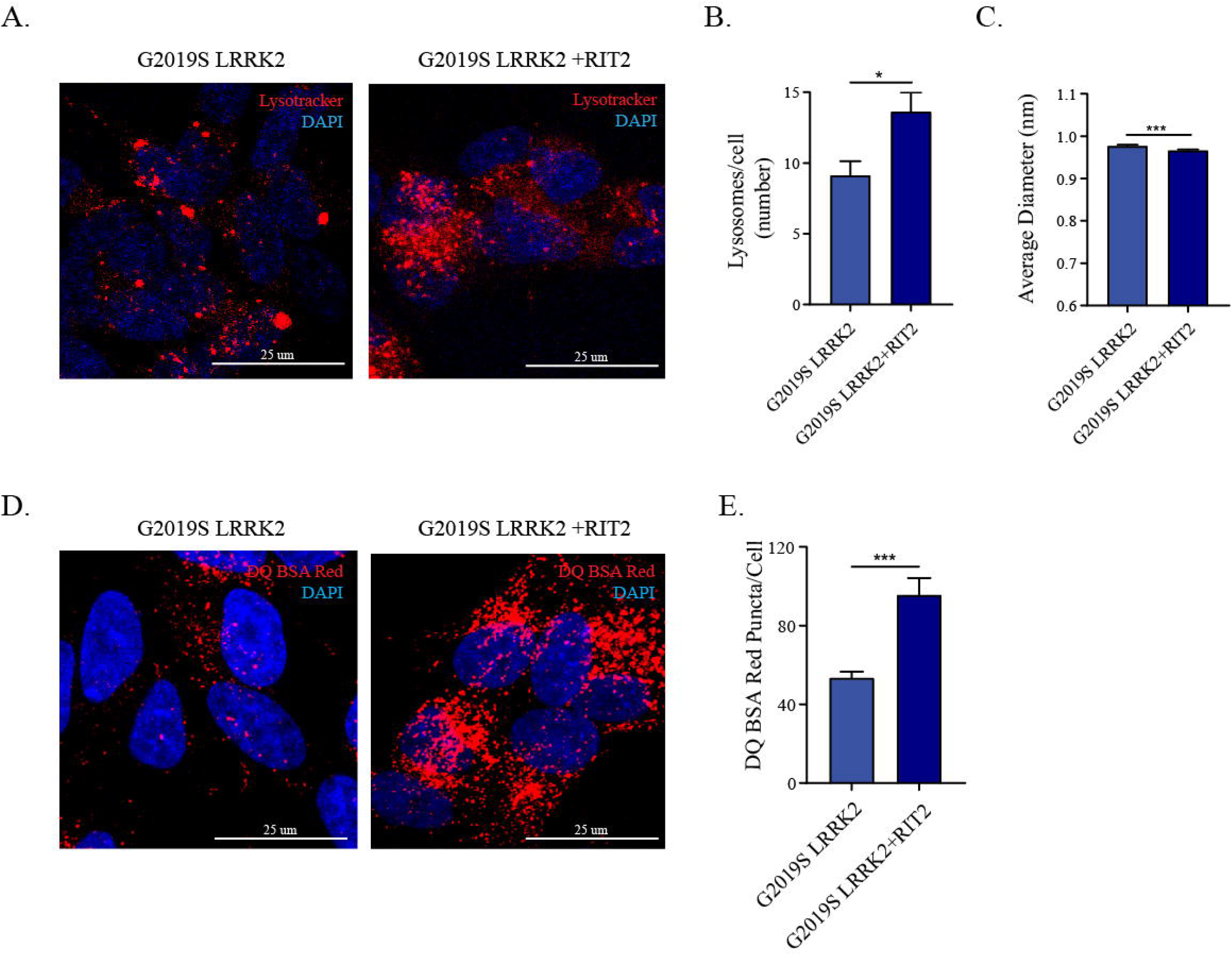
RIT2 overexpression is rescuing lysosomal morphology and proteolytic activity in G2019S-LRRK2 cells. A) Cell processing with the Lysotracker Red dye was performed to visualize lysosomes G2019S-LRRK2 and G2019S LRRK2+*RIT2* cells. B) The number of lysosomes per cell was quantified and revealed an increase, when *RIT2* was transfected in G2019S LRRK2 cells (n=3). C) The average size of lysosomes was assessed, and a significant decrease of the diameter was measured when *RIT2* was transfected to G2019S-LRRK2 cells (n=3). D) The DQ-Red-BSA assay was employed to assess the proteolytic activity of lysosomes. E) Quantification of DQ-Red-BSA fluorescent spots revealed a significant increase in G2019S LRRK2 cells, with *RIT2* overexpression (n=3). In imaging experiments, 3 independent biological replicates were performed and analysis conducted on 700-1000 cells per group in each experiment. *p<0.05, ***p<0.001, unpaired two tailed Student’s t-test.

**Figure 4:**
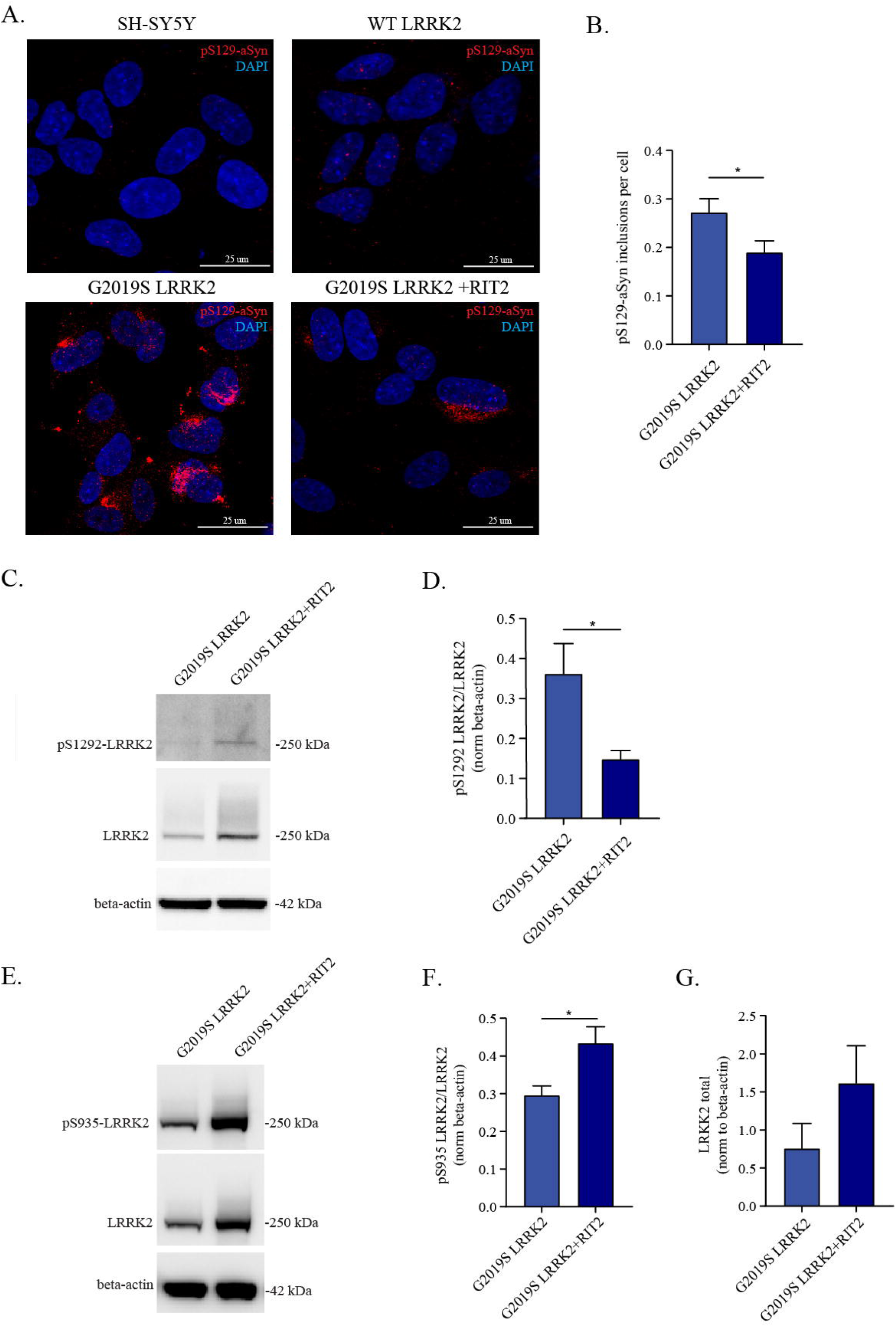
RIT2 overexpression reduces pS129-aSyn positive inclusions and reduces pS1292-LRRK2, but not pS935 LRRK2 levels. A) Representative images showing pS129-aSyn immunostaining in SH-SY5Y, WT-LRRK2 and G2019S-LRRK2 cells with or without *RIT2* overexpression. B) Quantification pS129-aSyn in cell overexpressing either G2019S-LRRK2 alone or G2019S-LRRK2 with *RIT2*. Four independent biological replicates were performed and analysis was conducted on 700-1000 cells per group in each experiment. C) Phosphorylation levels of S1292 in G2019S-LRRK2 and G2019S LRRK2+*RIT2* were measured using WB for pS1292-LRRK2 and total LRRK2. D) pS1292 LRRK2 levels are reduced when *RIT2* is overexpressed (normalized to total LRRK2) (n=5). E) Phosphorylation levels of S935 in G2019S-LRRK2 and G2019S LRRK2+RIT2 were measured using WB for pS935-LRRK2 and total LRRK2 F) pS935 LRRK2 levels are increased when *RIT2* is overexpressed and when normalized to total LRRK2 (n=6). G) *RIT2* overexpression leads to a trend of increased total LRRK2 levels, when normalized to b-actin. Data are means±SEM of 5-6 independent experiments for WB. *p<0.05, unpaired two tailed Student’s t-test.

**Figure 5:**
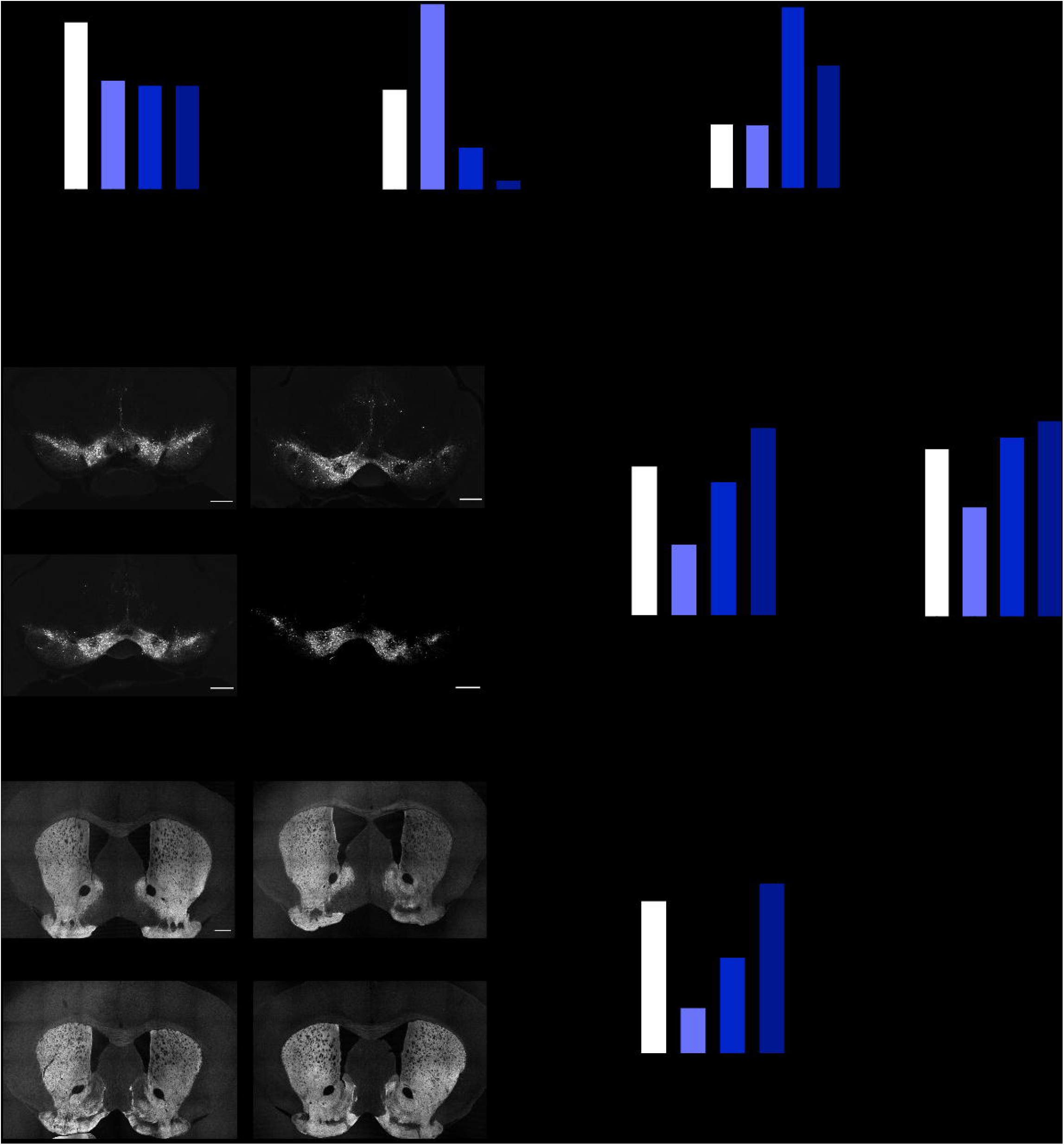
Enhanced RIT2 expression in the mouse midbrain counteracts aSyn-dependent deficits and DA neuron loss. A) Overexpression of A53T-aSyn reduces the percentage of contralateral forepaw use and co-injection with AAV-Flex-*Rit2* has the same effect. B) Overexpression of A53T-aSyn increases the number of ipsilateral rotations and co-injection with AAV-*Rit2* or injection of AAV-*Rit2* alone increases the number of contralateral rotations. C) Overexpression of *Rit2* alone or with aSyn significantly increases horizontal activity in the open field (n^GFP^=7, n^aSyn^=9, n^aSyn+RIT2^=8, n^RIT2^=6 for behavioural tests). D) IHC for TH was used to count dopaminergic neurons in the midbrain. E) Overexpression of A53T-aSyn alone induces a significant loss of DA neurons, which is attenuated by co-injection of AAV-Flex-*Rit2*. F) aSyn overexpression tends to reduce the number of NeuN+ cells in the ipsilateral SNc and concomitant overexpression of *Rit2* significantly preserves NeuN+ cells (5 animals/group). G) IHC for TH was used to measure the density of DA projections in the striatum. H) Overexpression of A53T-aSyn decreases the number of TH+ axons in the striatum, that is attenuated by the co-injection of AAV-Flex-*Rit2* (n=5/group for IHC experiments). Data represented as mean ± SEM *p<0,05 **p<0,01 ***p<0,001 ****p<0,0001, one-way ANOVA followed by Sidak’s test. Scale bar = 500 μm.

### Overexpression of RIT2 rescues ALP deficits in G2019S-LRRK2 cells

It has been extensively reported that both aSyn and LRRK2 affect the autophagic process (Bellomo et al., 2020; Bravo-San Pedro et al., 2013; Ho et al., 2019; Schapansky et al., 2014). Furthermore, we recently found that the reduction of pS129-aSyn-positive inclusions in G2019S-LRRK2 cells, following exposure to a LRRK2 kinase inhibitor, depends on correct lysosomal activity (Obergasteiger et al., 2020). As an unbiased approach we used the RT^2^ Profiler^™^ PCR Array for autophagy to measure the expression levels of 84 genes implicated at different stages in the ALP. Since G2019S-LRRK2 cells displayed lower *RIT2* mRNA levels, we decided to reconstitute *RIT2* expression in these cells, using high efficiency nucleofection of a construct containing the human *RIT2* gene (Fig. S2A) and investigate gene expression changes (Table S1). *RIT2* overexpression seemed to affect several genes related to autophagy; specifically, *WIPI1* and *DRAM1* gene expression levels were enhanced by *RIT2* overexpression. *WIPI1* promotes autophagy and autophagosome formation (Tsuyuki et al., 2014; Xiao et al., 2018), whereas *DRAM1* is implicated in lysosomal acidification, clearance of autophagosomes and fusion of autophagosomes with lysosomes (Zhang et al., 2013). Overall, the results obtained from the array suggested that the ALP is affected by the overexpression of *RIT2*. Therefore, we looked at specific stages of the ALP, starting with monitoring the overall autophagic flux.

G2019S-LRRK2 cells were transiently nucleofected with *RIT2*. After 72 hours, cellular extracts from G2019S-LRRK2 cells and G2019S-LRRK2+*RIT2* cells were prepared and the conversion of LC3B-I to LC3B-II was measured via Western blotting. Changes in the ratio of LC3B-II/LC3B-I are used to quantify the dynamics of the autophagic process. The LC3B-II/LC3B-I ratio was not significantly changed between G2019S-LRRK2 cells alone or transfected with *RIT2* (Fig. 2B), indicating that autophagy initiation is not affected by *RIT2* overexpression. In parallel, we assessed the LC3B-II/beta-actin ratio in the same cellular extracts, which was not affected by *RIT2* overexpression (Fig. 2C). Altogether, these data indicate that *RIT2* did not modulate the overall autophagic flux in G2019S-LRRK2 cells.

The Cyto-ID autophagy assay was used to visualize autophagosomes and the dye is also partially staining autolysosomes (Fig. 2D). *RIT2* overexpression increased the number of autophagosomes (Fig. 2E). We then evaluated the endpoint effector of the ALP, the lysosome. We investigated lysosomal morphology and number employing the Lysotracker assay (Fig. 3A), which accumulates in acidic compartments in the cell. We observed that *RIT2* overexpression increases the number of lysosomes (Fig. 3B) and decreases their size (Fig. 3C). Given these morphological effects, we measured the proteolytic function of lysosomes using the DQ-Red-BSA assay. This artificial BSA is degraded by proteases in the acidic environment of the lysosome and an increase of fluorescent spots indicates an enhanced amount of cleaved DQ-Red-BSA, and thus elevated lysosomal function. The overexpression of *RIT2* significantly enhanced the number of fluorescent spots per cell (Figure. 3D-E), indicating an increase in lysosomal proteolysis. Taken together these results reveal that *RIT2* does not affect autophagy initiation or the overall autophagic flux, but increases autophagosome and lysosome number, likely through the rescue of morphological and functional deficits of the lysosomes.

### pS129-aSyn positive inclusions in G2019S-LRRK2 cells are reduced by RIT2 overexpression

A significant proportion of PD patients carrying the G2019S mutation exhibit LB pathology (Henderson et al., 2019; Kalia et al., 2015) and, in experimental models, this mutation has been shown to promote the accumulation of pS129-aSyn (Volpicelli-Daley et al., 2016). SH-SY5Y neuroblastoma cell lines stably overexpressing G2019S-LRRK2 display inclusion-like structures staining positively for pS129-aSyn, while WT-LRRK2 cells only exhibited pale, diffuse pS129-aSyn staining comparable to naïve cells (Fig. 4A). Enhanced expression of *RIT2* showed beneficial effects on different stages of the ALP; therefore, we hypothesized it might play a role in the degradation of pS129-aSyn. Thus, we decided to investigate the effects of *RIT2* on pathological inclusions. G2019S-LRRK2 cells were transiently nucleofected with *RIT2* or GFP control and after 72 hours of overexpression, cells were fixed and processed for pS129-aSyn staining. We investigated the number of pS129-aSyn-positive objects per cell. Overexpression of *RIT2* significantly reduced the number of objects positively stained for pS129-aSyn per cell (Fig. 4A-B). Control GFP nucleofection did not alter pS129-aSyn inclusion number (Fig. S2B-C).

### RIT2 overexpression reduces kinase activity of recombinant LRRK2

Our results indicate that *RIT2* overexpression in G2019S-LRRK2 cells phenocopies pharmacological LRRK2 kinase inhibition as we recently reported (Obergasteiger et al., 2020). Thus, we hypothesized that LRRK2 and *RIT2* could play a role in a common molecular mechanism. We explored the possibility that LRRK2 kinase inhibition could influence *RIT2* mRNA levels, with LRRK2 acting upstream of *RIT2*. We treated G2019S-LRRK2 cells with different concentrations of the LRRK2-selective kinase inhibitor PF-06447475 (Daher et al., 2015) and *RIT2* mRNA levels were assessed using ddPCR. No difference in *RIT2* gene expression levels were observed upon pharmacological LRRK2 kinase inhibition (Fig. S3). Since LRRK2 kinase inhibition did not affect *RIT2* expression, we explored the opposite hypothesis, i.e. that *RIT2* overexpression inhibits LRRK2 kinase activity (*RIT2* acting upstream of LRRK2). LRRK2 phosphorylation was measured using Western blotting at two distinct residues (S935 and S1292), which are responsive to pharmacological LRRK2 inhibition (Fig. 5) (Kluss et al., 2018). The autophosphorylation of S1292 (pS1292) is a validated readout of LRRK2 kinase activity (Kluss et al., 2018). We previously found that G2019S-LRRK2 cells harbor a 3-4-fold increase of relative pS1292-LRRK2 when compared to WT-LRRK2 cells (Obergasteiger et al., 2020). The measurement of LRRK2 phosphorylation following overexpression of *RIT2* revealed a significant decrease of pS1292-LRRK2 levels (Fig. 4C-D). Differently from LRRK2 kinase inhibitor treatment, pS935-LRRK2 levels were not decreased, but rather significantly increased (Fig. 4E-F). Moreover, total LRRK2 levels in G2019S-LRRK2 cells overexpressing *RIT2* were slightly increased (Fig. 4G). This effect is distinct from LRRK2 kinase inhibitors, which have been shown to reduce total LRRK2 through protein destabilization (Lobbestael et al., 2016). Consistent with a lack of S935 dephosphorylation, our results indicate that *RIT2* does not reduce LRRK2 protein stability.

### Enhanced Rit2 expression in the mouse midbrain counteracts aSyn-dependent deficits and DA neuron loss

To determine if the beneficial effects of *RIT2* could also be translated *in vivo*, we modeled PD pathology in mice by unilaterally injecting an Adeno-Associated virus 2/9 (AAV) expressing the human A53T-aSyn (AAV-CMVie-hSynP-synA53T, herein AAV-A53T-aSyn). We used DAT-Ires-Cre mice that specifically express the Cre recombinase in neurons expressing the dopamine transporter (DAT) (Backman et al., 2006). This allowed us to manipulate gene expression in a neuron-specific manner, as *RIT2* is specifically downregulated in DA neurons in PD patients (Fig. 1A). The AAV-mediated overexpression of aSyn in the mouse midbrain has been previously shown to induce progressive SNc neuron loss, aSyn pathology and motor deficits (Oliveras-Salva et al., 2013). Sixteen weeks following overexpression of A53T-aSyn (in combination with AAV-CAG-Flex-EGFP), we observed motor abnormalities in the cylinder, rotation and amphetamine tests (Fig. 5A-B and Fig. S4A) with an ipsilateral rotation phenotype and a preferential use of the ipsilateral forepaw. In addition, A53T-aSyn overexpression induced a loss of TH-positive neurons in the midbrain (Fig. 5D-E) coupled to a loss of striatal DA terminals (Fig. 5G-H), mimicking the progressive nigrostriatal degeneration observed in PD. Overexpression of A53T-aSyn also tended to decrease the number of NeuN-positive cells in the ipsilateral SNc (Fig 5F). To further ensure that overexpression of A53T-aSyn did not cause a downregulation of TH expression without cell loss, we counted the percentage of double-positive cells for GFP and TH in the ipsilateral SNc of AAV-GFP and AAV-GFP+A53T-aSyn injected mice. Both groups showed a percentage of double-positive cells of around 90%, suggesting that A53T-aSyn overexpression did not alter TH expression in dopaminergic neurons (Fig. S4). Mice were co-injected with AAV-A53T-aSyn and the Cre dependent AAV-CAG-Flex-*Rit2*-EGFP (AAV-Flex-Rit2) or AAV-CAG-Flex-EGFP (AAV-GFP) control, to achieve overexpression selectively in DA neurons. This strategy ensured that the observed phenotypes would be attributed to the effect of *Rit2* in this neuronal population. The loss of TH-positive neurons in the SNc (Fig. 5E) and of DA terminals in the striatum (Fig 5H) were greatly attenuated by the co-expression of *Rit2*, when compared to GFP control, and also significantly preserved NeuN+ cells in the ipsilateral SNc (Fig 5F). Overexpression of *Rit2* alone did not cause loss of midbrain neurons nor striatal terminals, but interestingly, strongly promoted locomotor activity. We observed a marked increase in horizontal activity (Fig. 5C) in the open field, along with an increased number of contralateral rotations in the cylinder test (Fig. 5B).

### Enhanced RIT2 expression reduces pS129-aSyn load and total aSyn levels

Viral overexpression of A53T-aSyn in the mouse SNc also led to a significant increase of total aSyn protein levels measured by Western blotting (Fig. 6A-B). The concomitant overexpression of *Rit2* produced a reduction of total aSyn, when compared to AAV-GFP coinjection (Fig. 6A-B). Importantly, viral A53T-aSyn overexpression significantly increased the load of pS129-aSyn in DA neurons, when compared to AAV-GFP-injected animals. The co-injection of AAV-Flex-*Rit2* significantly reduced pS129-aSyn immunostaining in midbrain DA neurons (Fig 6C-D). This indicates that enhanced *Rit2* expression counteracts pS129-aSyn accumulation not only in recombinant neuroblastoma cell lines, but importantly also in an *in vivo* PD mouse model.

**Figure 6:**
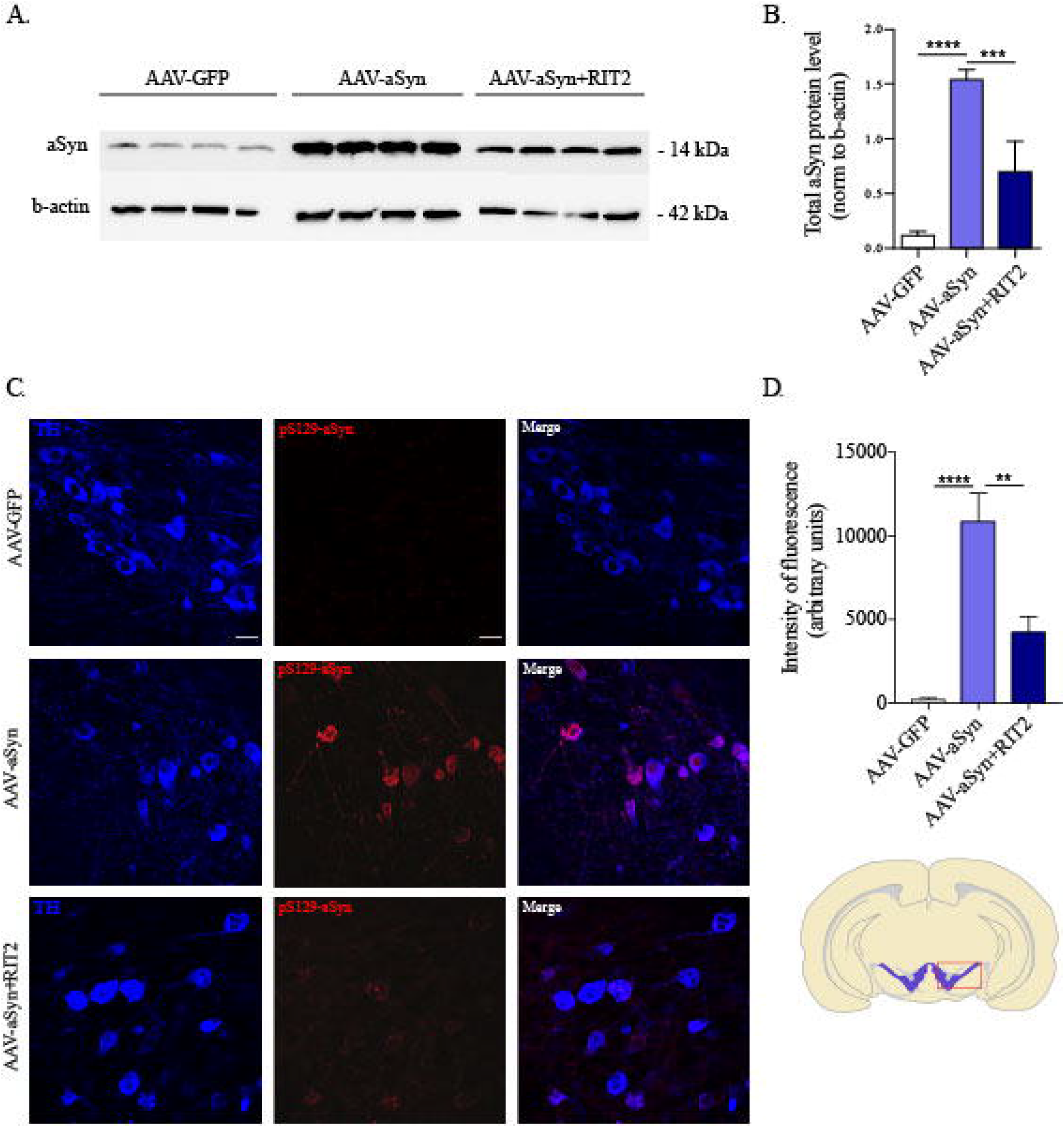
Enhanced RIT2 expression reduces total aSyn and pS129-aSyn levels. A) Total aSyn levels in mice injected with AAV-GFP, AAV-A53T-aSyn alone or in combination with AAV-Flex-*Rit2* were assessed blotting for total aSyn and β-actin. B) Quantification of total aSyn levels, normalized on β-actin (4 animals/group). Co-injection of *AAV-Flex-Rit2* significantly reduces aSyn levels in the ipsilateral side. C) IHC staining of pS129-aSyn and TH in the midbrain of AAV GFP, AAV-A53T-aSyn and AAV-A53T-aSyn + AAV-Flex-*Rit2* injected mice. D) Quantification of pS129-aSyn intensity. AAV-A53T-aSyn injection significantly increases the intensity of pS129-aSyn signal, which is reduced by the co-injection of AAV-Flex-*Rit2* (5 animals/group). Data represented as mean ± SEM. **p<0.01, ***p<0.001, ****p<0.0001, one-way ANOVA followed by the Bonferroni’s post-hoc test. Scale bar = 20 μm.

### RIT2 overexpression prevents aSyn-induced endogenous LRRK2 kinase hyperactivation

We show that viral co-expression of *Rit2* with A53T-aSyn in the mouse rescues neuronal loss and ameliorates behavioural deficits. In a recent study, viral overexpression of aSyn in the rat leads to increased kinase activity of endogenous LRRK2, observed as well in the midbrain of idiopathic PD patients, in the absence of any LRRK2 mutation (Di Maio et al., 2018). *RIT2* reduces phosphorylation levels of S1292-LRRK2 in recombinant neuroblastoma cells, therefore we assessed the impact of *Rit2* on S1292 phosphorylation of endogenous, murine LRRK2 in the mouse DA neurons. Proximity ligation assay (PLA) was employed on mouse midbrain sections (Fig. 7A), because it increases specificity and decreases background with respect to standard immunostaining (Di Maio et al., 2018). The injection of AAV-A53T-aSyn induced a strong increase in the number of PLA dots, indicating a significant enhancement of endogenous LRRK2 phosphorylation at S1292 and thus an increase in kinase activity. *Rit2* co-expression completely prevented the overactivation of endogenous LRRK2 that was elicited by A53T-aSyn (Fig. 7B). Importantly, total LRRK2 levels were not changed in any of the conditions analyzed (Fig. S5). In summary, viral *Rit2* overexpression in the mouse SNc neurons conferred neuroprotection, reduced aSyn neuropathology through prevention of endogenous LRRK2 hyperactivation and rescued A53T-aSyn-induced motor deficits.

**Figure 7:**
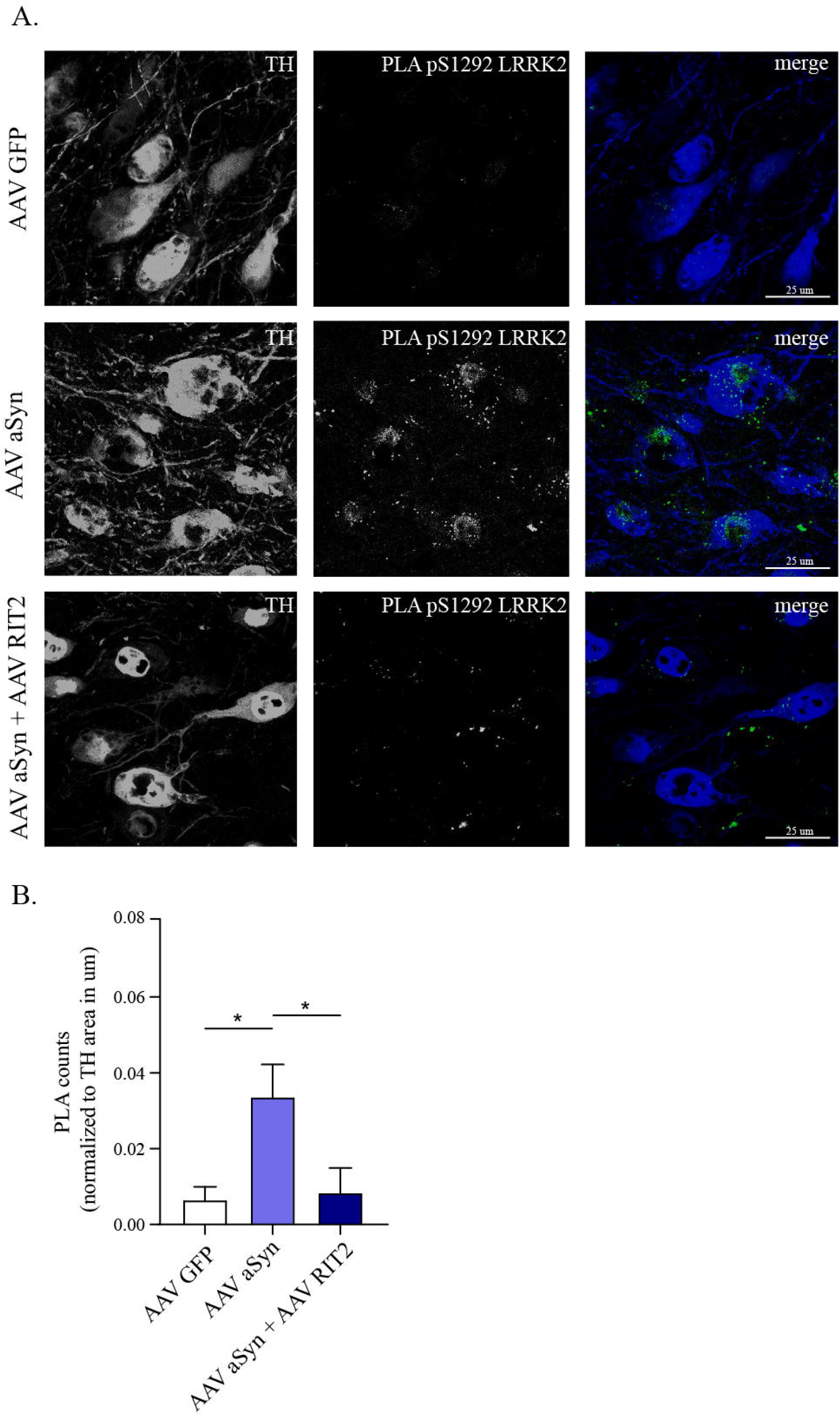
In vivo aSyn overexpression increases endogenous LRRK2 activity, which is prevented by RIT2 co-expression. A) PLA analysis of AAV-GFP, AAV-A53T-aSyn and AAV-A53T-aSyn+AAV-Flex-*Rit2* injected mice in TH-positive neurons in the SNc. B) Quantification of PLA counts in TH-positive neurons shows a significant increase of endogenous LRRK2 kinase activity with AAV-A53T-aSyn injection. The increase is completely prevented by co-injection of AAV-Flex-*Rit2* with AAV-A53T-aSyn (AAV-GFP=6 animals, AAV-A53T-aSyn=6 animals, AAV-A53T-aSyn+AAV-Flex-*Rit2*=5 animals). *p<0.05, one-way ANOVA followed by Bonferroni’s post-hoc test.

## Discussion

In this study, we aimed at evaluating the potential neuroprotective role of a novel, underexplored target against aSyn neuropathology. We validated the potential of the small GTPase *RIT2* in complementary *in vitro* and *in vivo* models of PD and revealed the cellular mechanism involved. In recent GWAS, *RIT2* has been associated to PD (Pankratz et al., 2012), and neuropsychiatric disorders (Glessner et al., 2010; Hamedani et al., 2017; Liu et al., 2016). Here, we show *RIT2* mRNA levels are reduced in DA neurons from idiopathic PD patients, in G2019S-LRRK2 overexpressing cells and mouse midbrain after AAV-A53T aSyn expression. Altogether, expression analysis shows that *RIT2* is expressed by DA neurons and reduced in PD conditions.

The downregulation of *RIT2* mRNA levels constitutes the rationale to transiently overexpress *RIT2* in G2019S-LRRK2 cells. *RIT2* overexpression was effective in reducing autophagosome number, increasing lysosome number and reducing their size. Moreover, *RIT2* was able to rescue the impaired proteolytic function of lysosomes. In summary, enhanced *RIT2* expression provided similar effects to those observed with the application of a LRRK2 kinase inhibitor.

Consistently, the overexpression of *RIT2* reduced the number of pS129-aSyn inclusions in G2019S-LRRK2 cells. It has already been reported that autophagy impairment might constitute a causative factor in aSyn accumulation (Cuervo et al., 2004; Komatsu et al., 2006; Lynch-Day et al., 2012) and autophagy/lysosome markers are altered in PD brain areas affected by LB pathology (Chu et al., 2009; Dehay et al., 2010). In our study, we specifically focused on macroautophagy since, in our cell model, this major form of autophagy was found to be affected. Nevertheless, we do not exclude an involvement of CMA that is also profoundly implicated in LRRK2 biology and aSyn pathology (Cuervo et al., 2004; Ho et al., 2019; Orenstein et al., 2013; Webb et al., 2003). Notably, the impairment of (macro)autophagy leads to an activation of CMA and vice versa, in a bi-directional compensatory mechanism (Kaushik et al., 2008).

Our results suggest that *RIT2* and LRRK2 function in a common pathway. We found that *in vitro RIT2* overexpression modulates the phosphorylation levels of LRRK2. Importantly, S1292 phosphorylation was reduced by *RIT2* overexpression, whereas pS935-LRRK2 was significantly increased, critically differentiating *RIT2*-related effects from those of pharmacological LRRK2 kinase inhibition (which targets both phosphosites). The G2019S-LRRK2 mutation is known to increase aSyn inclusion formation after application of aSyn preformed fibrils. Indeed, it has been reported that LRRK2 kinase inhibition is beneficial against aSyn aggregation and neurodegeneration, highlighting that these effects are dependent on LRRK2 kinase activity (Volpicelli-Daley et al., 2016) and even more important because PD-associated mutations in LRRK2 increase its kinase activity (West et al., 2005). Similar beneficial effects on aSyn inclusion formation and neuron loss were achieved by reducing total LRRK2 levels using antisense oligonucleotides against LRRK2 (Zhao et al., 2017), strengthening the important role of LRRK2 kinase activity in aSyn aggregation and associated to defective autophagy. In addition we show that total LRRK2 protein levels were not reduced when overexpressing *RIT2*, contrasting with the effect of LRRK2 kinase inhibitors (Lobbestael et al., 2016). Phosphorylation of S935-LRRK2 is required for a stable binding to 14-3-3 proteins and a reduction in pS935-LRRK2 was shown to reduce this interaction. The disrupted interaction leads to mislocalization of LRRK2 into cytoplasmic pools and therefore, the S935-dependent 14-3-3 binding is thought to protect LRRK2 protein from degradation (Dzamko et al., 2010; Nichols et al., 2010). The exact mechanism behind LRRK2 destabilization is not yet understood, but it is believed to rely on LRRK2 kinase activity, since mice expressing a kinase-dead mutation in LRRK2 display reduced LRRK2 protein levels (Herzig et al., 2011; Mercatelli et al., 2019). Understanding the exact regulation of LRRK2 protein degradation is a key point to predict side effects of LRRK2 kinase inhibitors, as it has been reported to produce lung and kidney pathology, similar to observations in LRRK2 KO mice (Tong et al., 2010). LRRK2 inhibitors are, in fact, envisaged to be administered not only to LRRK2-PD patients, but also to idiopathic PD patients, as indicated by the increase of pS1292-LRRK2 levels in urinary exosomes and brain tissue (Di Maio et al., 2018; Fraser et al., 2016). We found that *RIT2* overexpression is capable of reducing pS1292-LRRK2 levels, without affecting S935-LRRK2 phosphorylation and total protein stability. These findings indicate that a potential strategy based on targeting *RIT2* or its associated pathway(s) could be efficient in inhibiting LRRK2 while avoiding side effects induced by LRRK2 kinase inhibitors and related to reduction of total protein levels. Notably, targeting *RIT2* could be a direct way to impact autophagy and aSyn aggregation.

Accordingly, our results indicate that *Rit2* overexpression not only rescued motor impairments, but also strongly attenuated the loss of DA neurons and striatal DA terminals. Moreover, total aSyn levels and pS129-aSyn pathology were reduced by *Rit2*. Our results suggest that enhanced *RIT2* expression has also beneficial effects on aSyn–inclusion formation.

Viral overexpression of *Rit2* in DA neurons, independently from aSyn overexpression, induced motor hyperactivity. This might be due to altered extracellular DA levels or release dynamics. It has been previously reported that RIT2 interacts directly with DAT, is required for DA trafficking and that RIT2 is necessary for the inactivation of DAT (Navaroli et al., 2011). This suggests that these modulations could be responsible for the increased locomotion. In fact, the genetic deletion of DAT in rats increases extracellular DA lifetime, resulting in locomotor hyperactivity (Leo et al., 2018). Interestingly, the conditional KO of *Rit2* in mouse midbrain DA neurons decreases total DAT protein in the striatum, strengthening the possible modulation of DAT by *RIT2* (Sweeney et al., 2019).

Lastly, the reduction of pS1292-LRRK2 levels *in vitro* and the recent observation that virally overexpressed aSyn in rats increases endogenous LRRK2 kinase activity (Di Maio et al., 2018), led us to investigate LRRK2 kinase activity in our mouse model. Using a viral vector encoding A53T-aSyn we confirmed the increased pS1292-LRRK2 levels in DA neurons, indicating an increased kinase activity. The co-expression of *Rit2* completely prevented the increase of pS1292-LRRK2 levels. As previously reported, aSyn overexpression results in an increase of endogenous LRRK2 activity, impacting the ALP and worsening aSyn pathology (Di Maio et al., 2018). These results suggest that the activation of wild-type LRRK2 plays a role in idiopathic PD pathogenesis and consequently holds a strong pathogenic implication in most PD cases.

However, upstream modifiers of LRRK2 and related mechanisms are mostly unknown. *Rab7L1*, a PD risk factor (Satake et al., 2009), has recently been suggested to recruit LRRK2 to the Golgi-network (Madero-Perez et al., 2018), stressed lysosomes (Eguchi et al., 2018) and to stimulate its kinase activity (Purlyte et al., 2018). Interestingly, Rab7L1 is a Rab GTPase, part of the RAS superfamily (Wang et al., 2014) to which RIT2 also belongs (Lee et al., 1996). Moreover, we previously proposed a novel hypothesis, in which GTPase-MAPK signaling is involved in the regulation of autophagy (Obergasteiger et al., 2018). Our study provides evidence for a molecular mechanism leading to LRRK2 kinase inhibition, with *RIT2* acting as an upstream modifier of LRRK2. This could partly explain the reduced penetrance of the G2019S-LRRK2 mutation, when considering possible differential *RIT2* expression.

In conclusion, we demonstrate that *RIT2* acts both on autophagy-related processes and pS129-aSyn clearance. Our results suggest that inhibiting LRRK2 kinase activity through enhanced RIT2 expression is beneficial against ALP defects, aSyn pathology and could target not only G2019S-LRRK2 PD, but also idiopathic PD. Our results suggest *RIT2*, through modulation of LRRK2 activity, as a novel target for neuroprotection in PD.

## Methods

### Cell culture and transfection

SH-SY5Y neuroblastoma cell lines stably overexpressing wild-type (WT) or G2019S-LRRK2 were previously described (Obergasteiger et al., 2017; Vancraenenbroeck et al., 2014) and maintained as in (Obergasteiger et al., 2020). SK-N-SH neuroblastoma cell lines stably overexpressing A35T-aSyn were recently described and maintained as in (Volta et al., 2017).

Cells were transfected using the SF Cell Line 4D-Nucleofector™ X kit (Lonza) with the 4D Nucleofector X unit. Nucleofection was carried out according to the manufacturers protocol. Briefly, 200.000 cells were resuspended in the nucleofection solution containing 800 ng plasmid DNA (SC118279, Origene, or eGFP in pcDNA3.1) and analysed after 72 h.

### Droplet digital PCR and autophagy qPCR array

Total RNA was extracted using the RNeasy Plus Mini Kit (Qiagen). First strand cDNA was synthesized using the SuperScript VILO cDNA Synthesis Kit (Invitrogen). 1 ng of cDNA was used for each reaction to asses *RIT2* mRNA gene expression (Hs01046671_m1, FAM), multiplexed with a housekeeping gene (RPP30, HEX, BioRad cat# 10031243). PCR was carried out as suggested by the manufacturer using ddPCR Supermix for probes (BioRad, cat# 11969064001).

PCR program

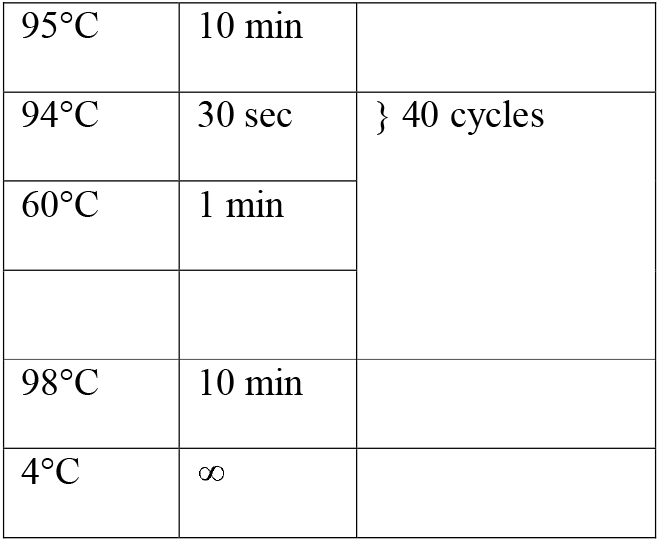

After PCR amplification, the plate was analyzed in the QX200 Droplet Reader. Data was analyzed using QuantaSoftTM software from BioRad. Results are reported as Fractional abundance of gene of interest respect to housekeeping gene.

The autophagy gene expression array (PAHS-084Z, Qiagen) for G2019S-LRRK2 cells without and with RIT2 overexpression was carried out according to the manufacturers protocol and as described in (Obergasteiger et al., 2020). Quantitative PCR was performed in a CFX96 Touch™ Real-Time PCR Detection System (BioRad). Analysis was carried out with the webtool provided by Qiagen (https://www.qiagen.com/it/shop/genes-and-pathways/data-analysis-center-overview-page/).

### Analysis of RIT2 mRNA levels from publicly available datasets

Geo Dataset GSE2014 composed of SN tissues isolated by laser capture microdissection from sporadic PD patients and controls was analyzed with 206984_s_at probe for RIT2. Geo Dataset GSE46798 composed of human iPSC-derived DA neurons from A53T mutant and corrected control lines was analyzed with ILMN_1796673 probe for *RIT2*. Lastly, Geo Dataset GSE17542 composed of laser captured SNpc and VTA from TH-GFP mice was analyzed with 1448125_at probe for *RIT2*.

### Western blotting

Cell or tissue lysates were prepared as previously described in (Obergasteiger et al., 2020). Primary antibodies used: anti-LRRK2 1:20000 (Abcam, ab172378), anti-pS935-LRRK2 1:1000 (Abcam, ab172382), anti-pS1292-LRRK2 1:1000 (Abcam, ab206035), anti-LC3 1:1000 (Cell Signaling Technologies, 3868), anti-β-actin 1:6000 (Sigma Aldrich, A-5316), anti-aSyn 1:2000 (Abnova, MAB5383). Chemiluminescence images were acquired using Chemidoc Touch (BioRad) and relative band intensity levels were calculated using ImageLab software (BioRad).

### CYTO-ID, Lysotracker Deep Red and DQ-Red-BSA staining

To investigate autophagosome number we used the CYTO-ID^®^ Autophagy Detection Kit (Enzo, ENZ-51031) following the manufacturer’s instructions. Briefly, cells were incubated with CYTO-ID^®^ Green Detection Reagent for 30 min at 37°C. After washing, DAPI for live imaging (Invitrogen, R37605) was added and cells were visualized live on the confocal microscope in an environmental chamber. To investigate lysosome morphology, we utilized the Lysotracker Deep Red dye (Molecular Probes, L12492) following the manufacturer’s instructions, as in (Obergasteiger et al., 2020).

To study the proteolytic activity of lysosomes, the fluorescent DQ-Red-BSA dye (Molecular Probes, D12051) was used following manufacturer’s instructions, as in (Obergasteiger et al., 2020).

### Immunofluorescence and Immunohistochemistry

Cells were fixed in 4% PFA, permeabilized, blocked and incubated with primary and secondary antibodies, as described in (Obergasteiger et al., 2020). The primary antibodie was mouse anti-pS129-aSyn 1:2000 (Abcam, ab184674). The secondary antibodie was: Donkey anti-Mouse Secondary Antibody, Alexa Fluor 555 (A31570). Visualization was performed using a Leica SP8-X confocal laser scanning microscope system equipped with an oil immersion 63X objective and images were analysed using CellProfiler to quantify the number and intensity of investigated objects (object size in pixel units: pS129-aSyn 20-250; pipelines available upon request). Immunohistochemistry (IHC) was performed as described in (Salesse et al., 2020). For DA neuron survival quantification, sections of midbrain (4 sections interval) were stained with NeuN (Millipore, MAB377, 1:500) and TH (Pel-Freez Biologicals, P60101, 1:1000) antibodies, followed by incubation with Cy3 (Jackson Immuno, 715-165-150, 1:200) and Alexa Fluor 647 (Life Technologies, A-21448, 1:400) respectively. Images were acquired using a slide scanner (TissueScope^™^, Huron Digital Pathology). Both NeuN and TH positive neurons were counted using a stereology software with optical fractioning (Stereo Investigator, mbf bioscience). For fluorescence quantification of the striatal dopamine terminals, sections were labeled with a TH antibody (P60101) and incubated with Alexa Fluor 647 (Life Technologies, A-21448, 1:400). For aSyn neuropathology, sections were labeled for pS129-aSyn (*custom antibody*, 73C6 1μg/ml, see following section) followed by Alexa Fluor 647 (Life Technologies, A-31571, 1:400). We used our custom 73C6 antibody for pS129-aSyn quantification because the signal obtained on brain sections was stronger and more specific than with ab184674 (used on cells). Images were acquired with a confocal microscope (LSM700, Carl Zeiss) and fluorescence quantification was obtained by using Corrected Total Cell Fluorescence with ImageJ software.

### 73C6 antibody production and purification

Briefly, mice were immunized with a peptide containing amino acids 111 to 135 (C-terminal) from human aSyn. B-cells were harvested and fused with myeloma cells. Hybridomas were maintained in DMEM high glucose supplemented with 10% Ultra-low IgG Fetal Bovin Serum (FBS) and Penicillin/Streptomycin (Gibco) and filtered through a 0.2μm filter. For purification, Protein-G agarose (Thermo-Fisher, cat# P120399) was used according to manufacturer’s instructions. Final IgG concentration was determined with spectrometry (Nanodrop One, Thermo-Fisher). Antibody specificity was determined using Western Blotting on HEK293 cells transfected with either WT-aSyn or S129A-aSyn, with or without concomitant transfection with the PLK2 kinase (Fig. S6).

### Animals

Heterozygous DAT-Ires-Cre mice aged between two and three months from Jackson Laboratory were used (males and females). Housing, breeding and procedures were approved by the CPAUL (Comité de protection des animaux de l’Université Laval) and the CCPA (Comité canadien de protection des animaux).

### Stereotaxic injection of AAV vectors

Mice were deeply anesthetized with isoflurane (3-4% for induction and 2% for maintenance with 0.5% oxygen). AAV-emCBA-GFP-Flex (7.5E12), AAV-CMVie-hSynP-synA53T (7.5E12) + AAV-emCBA-GFP-Flex, AAV-CAG-Flex-*Rit2*-EGFP, AAV-CAG-Flex-*Rit2*-EGFP (7E12 GC/ml) + AAV-CMVie-hSynP-synA53T were unilaterally injected in the right substantia nigra. A total volume of 1ul was injected at 2nl/sec at the following coordinates: −3.5mm (A/P); +1mm (M/L); +4mm (D/V) from bregma. Mice were euthanized 4 months after surgery.

### Behavioral tests

#### Open field

Mice were placed in the room one hour prior to testing for habituation, and then were placed in the open field for 60 minutes. Open fields (**) were used in the 2-animal monitor mode and the activity of the mice was recorded automatically with the VersaMax software. The total distance traveled (cm), horizontal activity (number of beam breaks) and vertical activity (number of beam breaks) were analyzed. The test was performed after 1, 2, 3 and 4 months following injection.

### Cylinder, rotation and amphetamine test

Mice were placed, one hour prior to testing, in the room for habituation and were then placed in a cylinder with a diameter of 10 cm. Tests were performed 4 months after injection and were recorded by video. For the cylinder and rotation tests, mice were placed in the cylinder for 5 minutes. Forepaw use and complete rotations were analyzed manually by an investigator. For the amphetamine test, mice were injected with amphetamine at 5mg/kg of body weight. After 15 minutes, they were placed in the cylinder for 30 minutes. Their complete rotations were counted manually by an investigator.

### Fluorescence in situ hybridization

RNAscope probes for mouse TH and *RIT2* (TH – 317621, RIT2 – 58904) were designed by Advanced Cell Diagnostics (Newark, CA, USA). The *in situ* hybridization was carried out following the manufacturers protocol for fixed and frozen tissue and as described in (Salesse et al., 2020).

### Statistical analysis

Statistical analyses were performed using GraphPad Prism 8. One-way ANOVA was used in experiments comparing 3 or more groups, followed by Bonferroni’s or Sidak post-hoc test for pairwise comparisons. With 2 experimental groups, the unpaired two-tailed Student’s t-test or two-tailed Mann-Whitney test was utilized. Threshold for significance was set at p<0.05. All experiments were performed in a minimum of 3 independent biological replicates.

## Acknowledgments

The authors are thankful to: lab members Sara Pizzi and Giulia Lamonaca for their invaluable assistance and support; Dr. Francesca Pischedda and Prof. Giovanni Piccoli (University of Trento, Italy) for their help with the pS935-LRRK2 western blot; Anne Picard for the RT-PCR of autophagy genes; Dr. Alexandros Lavdas for the analysis of Lysotracker images. We thank Veronique Rioux for the technical assistance and Modesto Peralta for comments on the manuscript. We thank the CERVO Centre Molecular Tool Platform (https://neurophotonics.ca/fr/pom) for the production of the viral vectors used. The authors thank the Department of Innovation, Research and University of the Autonomous Province of Bozen/Bolzano for covering the Open Access publication costs.

## Author contributions

JO and AMC contributed to conceptualization, methodology, formal analysis, investigation, data curation, writing – original draft, writing – review and editing and visualization; GF contributed to methodology, formal analysis, investigation, data curation for some experiments; EL and VB contributed resources; CC contributed to the planning of some experiments and to revising the manuscript. AAH, PPP contributed to Eurac funding acquisition; CG contributed to methodology, formal analysis, investigation, data curation for some experiments. ML and MV contributed equally to conceptualization, methodology, formal analysis, data curation, writing – original draft, writing – review and editing, supervision, project administration, funding acquisition.

## Conflict of interest

The authors declare no competing interests.

## Funding

This work was supported by Parkinson Canada (grant number 2018-00157 to ML and MV), the Weston Brain Institute (grant number RR191071 to ML and MV), the Canadian Institute of Health Research to ML (420504), the Autonomous Province of Bozen/Bolzano for core funding to the Institute for Biomedicine of Eurac Research to MV. AMC received a scholarship from the Fonds de Recherche du Québec–Santé. ML receives salary support from Fonds de Recherche du Québec–Santé, Chercheur-Boursier Juniors 2 in partnership with Parkinson Québec (34974).

---

**Supplementary Table 1:** List of fold change of gene expression of autophagy array genes in G2019S LRRK2 NT and G2019S LRRK2 +RIT2 cells (normalized to G2019S LRRK2)

## Notes

### Competing Interest Statement

The authors have declared no competing interest.

### Summary of Updates

New data have been added in cell biology and in vivo experiments

