## Supplementary figures and images for "RIT2 reduces LRRK2 kinase activity and protects against alpha-synuclein neuropathology"

### Supplemental Figure 1

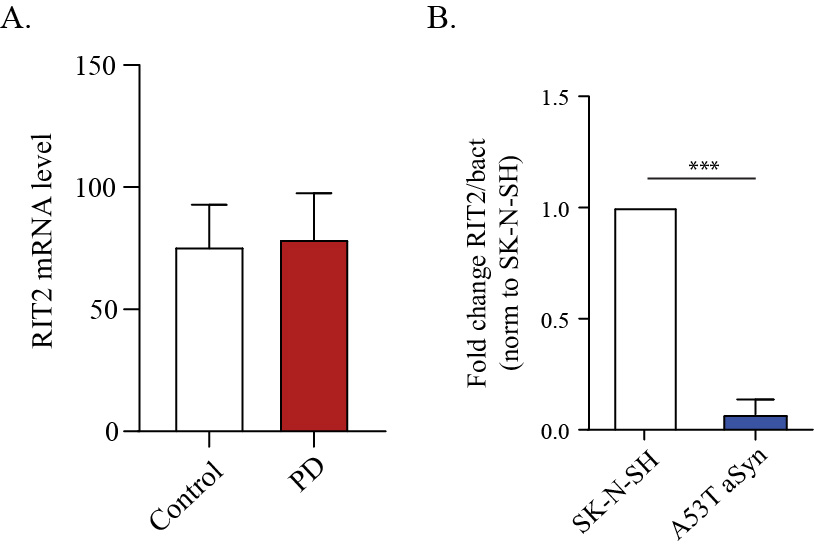

### Supplemental Figure 2

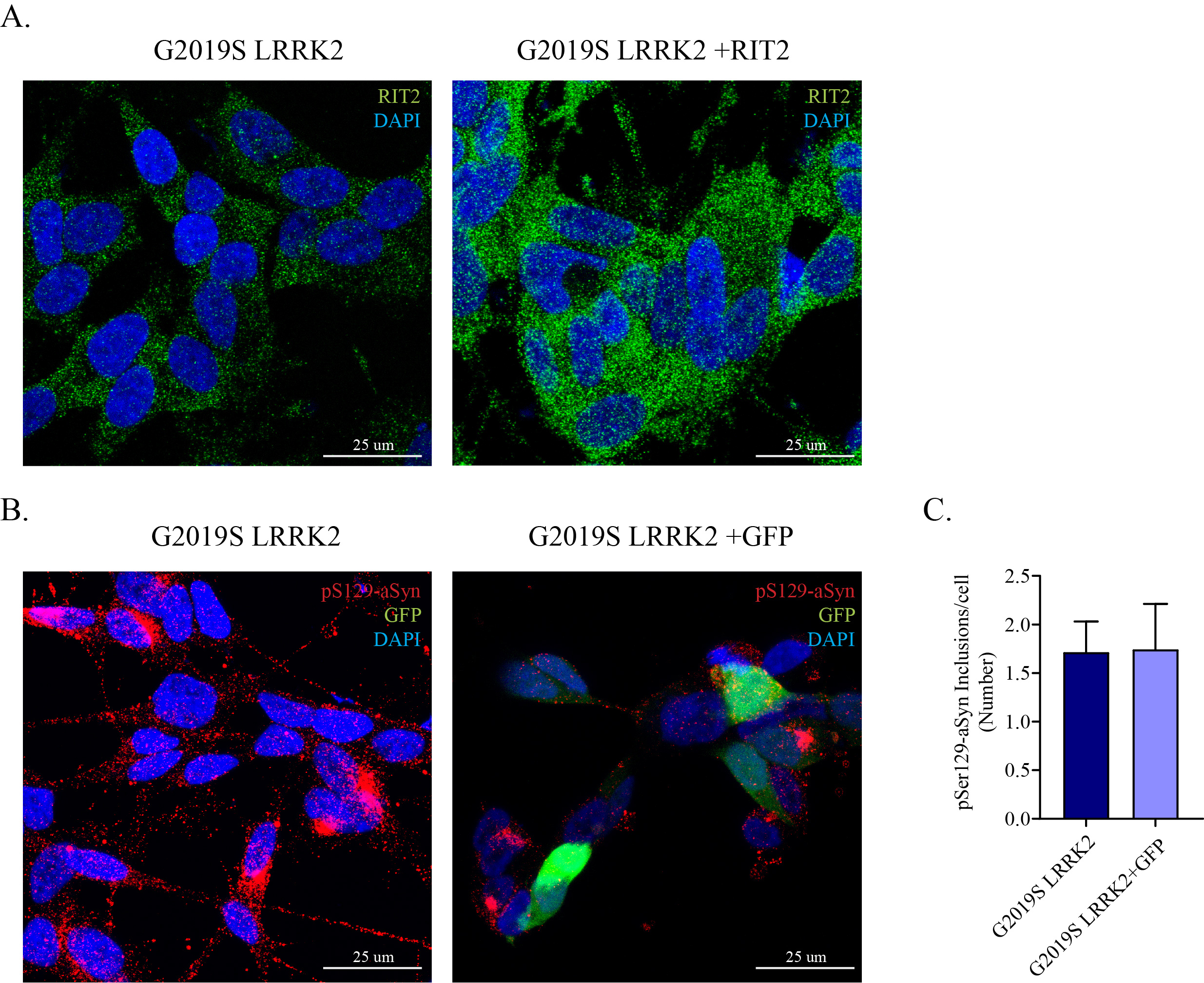

### Supplemental Figure 3

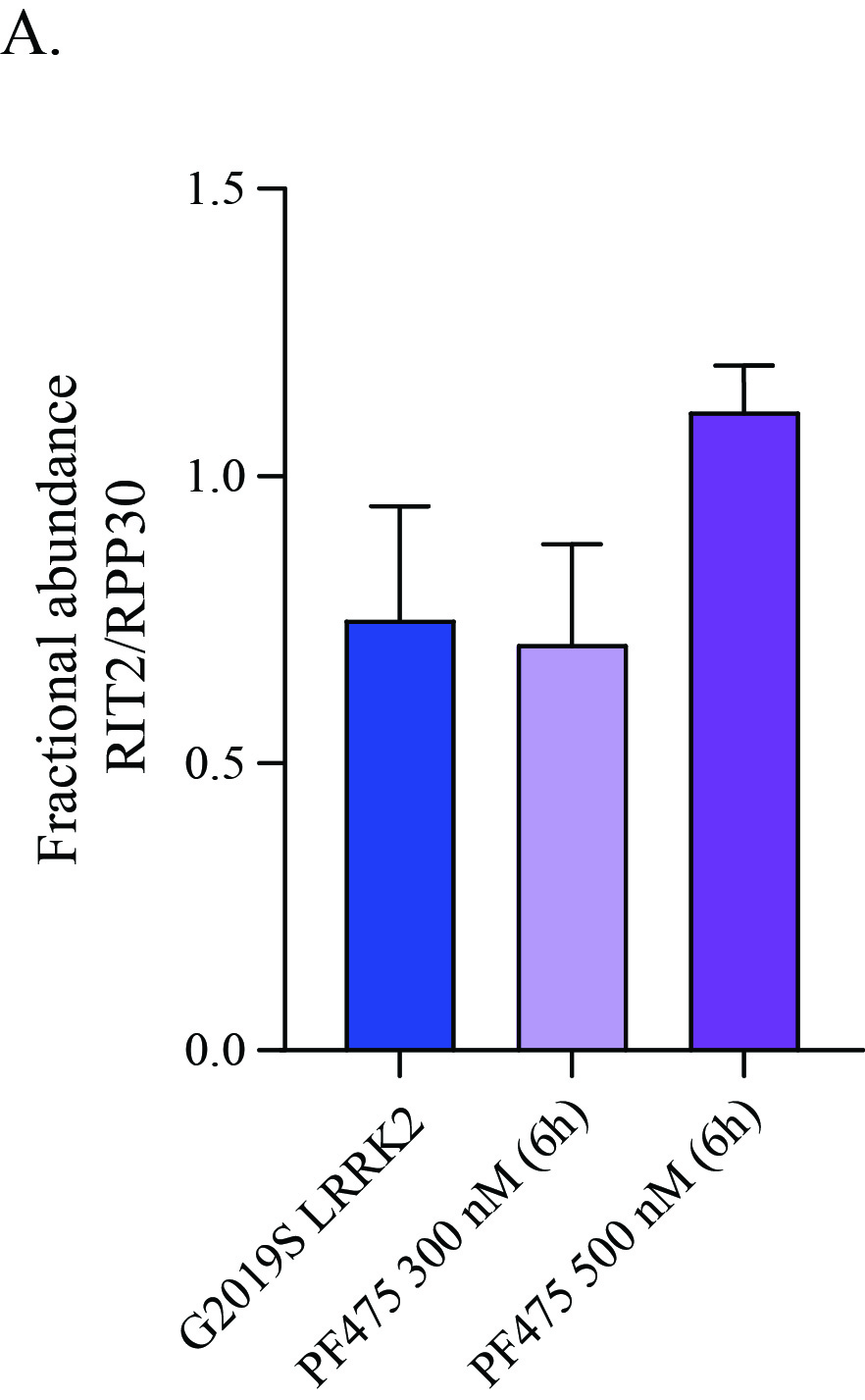

### Supplemental Figure 4

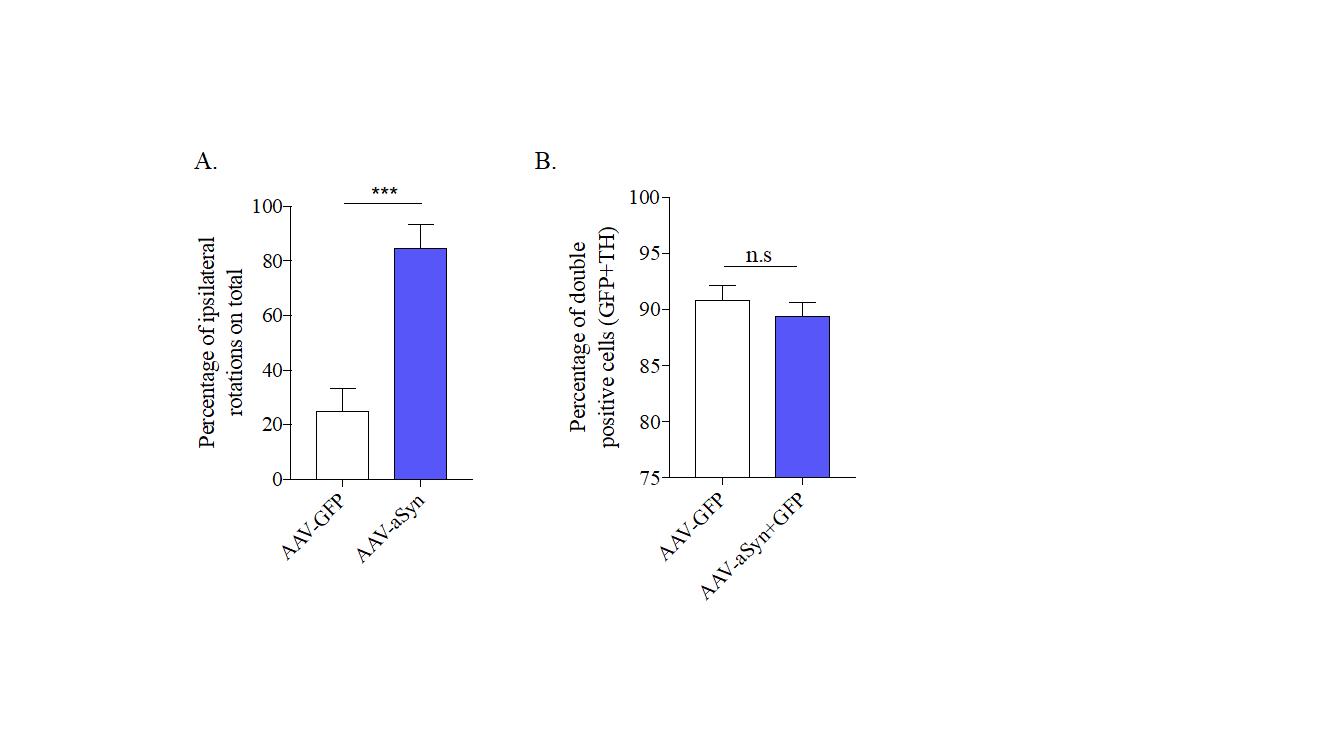

### Supplemental Figure 5

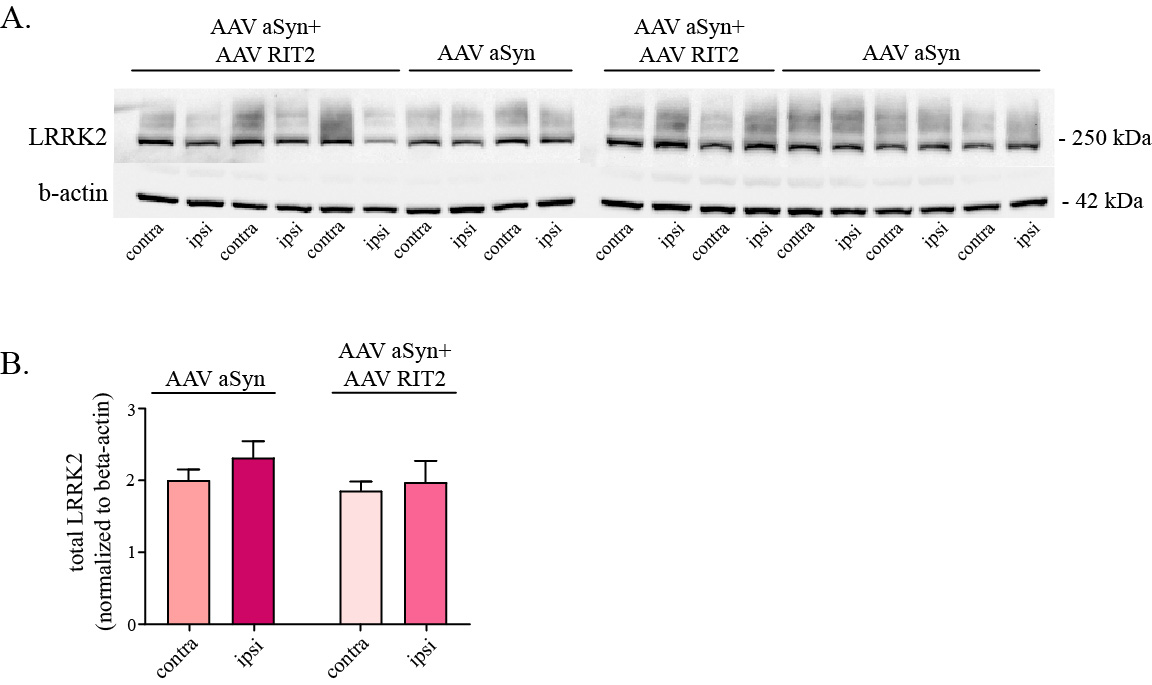

### Supplemental Figure 6

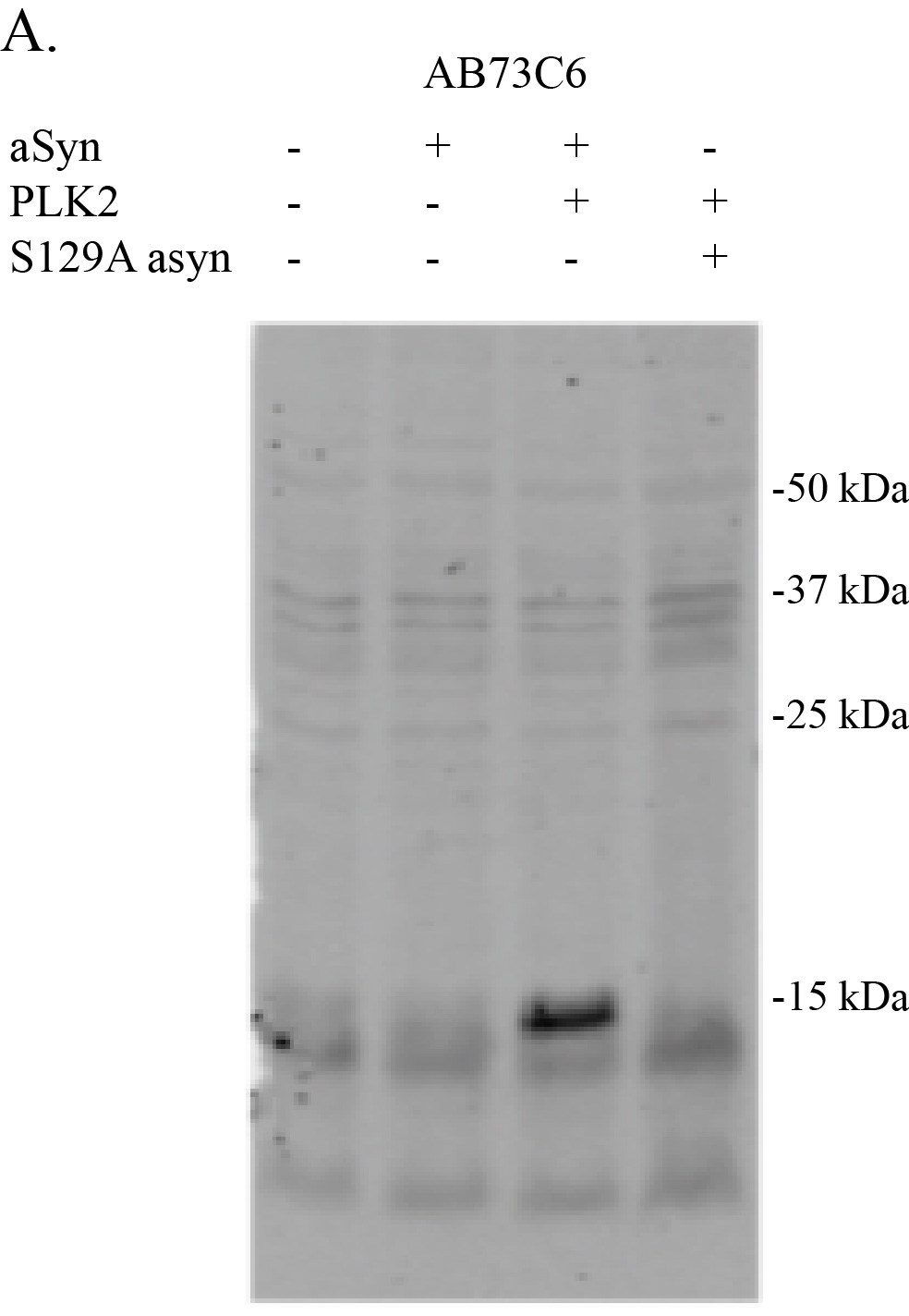
