## Supplemental Table 1 for "RIT2 reduces LRRK2 kinase activity and protects against alpha-synuclein neuropathology"

**Table EV1**: List of fold change of gene expression of autophagy array genes in G2019S LRRK2 NT and G2019S LRRK2 +RIT2 cells (normalized to G2019S LRRK2)

| **Gene symbol** | **G2019S NT** | **G2019S + RIT2** |
| --- | --- | --- |
| Autophagic Vacuole Formation: | | |
| AMBRA1 (NYW1) | 1,00 | 1,21 |
| ATG12 | 1,00 | 1,13 |
| ATG4A | 1,00 | 1,38 |
| ATG4B | 1,00 | 1,06 |
| ATG4C | 1,00 | 1,28 |
| ATG4D | 1,00 | 1,36 |
| ATG5 | 1,00 | 0,91 |
| ATG9A | 1,00 | 1,71 |
| ATG9B | 1,00 | 1,48 |
| BECN1 | 1,00 | 0,97 |
| GABARAP | 1,00 | 1,27 |
| GABARAPL2 | 1,00 | 0,95 |
| MAP1LC3A | 1,00 | 0,97 |
| MAP1LC3B | 1,00 | 0,92 |
| RGS19 | 1,00 | 1,14 |
| ULK1 | 1,00 | 1,20 |
| WIPI1 | 1,00 | 2,05 |
| Vacuole Targeting: | | |
| ATG4A | 1,00 | 1,38 |
| ATG4B | 1,00 | 1,06 |
| ATG4C | 1,00 | 1,28 |
| ATG4D | 1,00 | 1,36 |
| GABARAP | 1,00 | 1,27 |
| Protein Transport: | | |
| ATG10 | 1,00 | 1,21 |
| ATG16L2 | 1,00 | 1,71 |
| ATG3 | 1,00 | 0,94 |
| ATG4A | 1,00 | 1,38 |
| ATG4B | 1,00 | 1,06 |
| ATG4C | 1,00 | 1,28 |
| ATG4D | 1,00 | 1,36 |
| ATG7 | 1,00 | 0,94 |
| ATG9A | 1,00 | 1,71 |
| GABARAP | 1,00 | 1,27 |
| GABARAPL2 | 1,00 | 0,95 |
| RAB24 | 1,00 | 1,23 |
| Autophagosome-Lysosome Linkage: | | |
| DRAM1 | 1,00 | 2,36 |
| GABARAP | 1,00 | 1,27 |
| LAMP1 | 1,00 | 1,14 |
| NPC1 | 1,00 | 1,05 |
| Ubiquitination: | | |
| ATG3 | 1,00 | 0,94 |
| ATG7 | 1,00 | 0,94 |
| HDAC6 | 1,00 | 1,53 |
| Proteases: | | |
| ATG4A | 1,00 | 1,38 |
| ATG4B | 1,00 | 1,06 |
| ATG4C | 1,00 | 1,28 |
| ATG4D | 1,00 | 1,36 |
| Co-Regulators of Autophagy & Apoptosis: | | |
| AKT1 | 1,00 | 1,43 |
| APP | 1,00 | 1,09 |
| ATG12 | 1,00 | 1,13 |
| ATG5 | 1,00 | 0,91 |
| ATG7 | 1,00 | 0,94 |
| ATG9A | 1,00 | 1,71 |
| ATG9B | 1,00 | 1,48 |
| BAD | 1,00 | 1,44 |
| BAK1 | 1,00 | 0,96 |
| BAX | 1,00 | 1,37 |
| BCL2 | 1,00 | 1,07 |
| BCL2L1 | 1,00 | 0,99 |
| BECN1 | 1,00 | 0,97 |
| BID | 1,00 | 0,97 |
| BNIP3 | 1,00 | 1,28 |
| CASP3 | 1,00 | 0,92 |
| CDKN1B (P27KIP1) | 1,00 | 1,26 |
| CDKN2A (p16INK4a) | 1,00 | 1,11 |
| CLN3 | 1,00 | 1,19 |
| CTSB | 1,00 | 1,19 |
| CXCR4 | 1,00 | 1,63 |
| DAPK1 | 1,00 | 0,95 |
| DRAM1 | 1,00 | 2,36 |
| EIF1AK3 | 1,00 | 1,32 |
| FADD | 1,00 | 0,80 |
| FAS (TNFRSF6) | 1,00 | 1,19 |
| HDAC1 | 1,00 | 1,06 |
| HTT | 1,00 | 1,61 |
| MAPK8 (JNK1) | 1,00 | 0,96 |
| MTOR | 1,00 | 1,39 |
| NFKB1 | 1,00 | 1,20 |
| PRKAA1 (AMPK) | 1,00 | 1,20 |
| PTEN | 1,00 | 0,76 |
| SNCA | 1,00 | 1,18 |
| SQSTM1 | 1,00 | 1,18 |
| TGFB1 | 1,00 | 1,31 |
| TGM2 | 1,00 | 1,37 |
| TP53 (p53) | 1,00 | 1,22 |
| Co-Regulators of Autophagy & the Cell Cycle: | | |
| BAX | 1,00 | 1,37 |
| CDKN1B (P27KIP1) | 1,00 | 1,26 |
| CDKN2A (p16INK4a) | 1,00 | 1,11 |
| PTEN | 1,00 | 0,76 |
| RB1 | 1,00 | 1,19 |
| TGFB1 | 1,00 | 1,31 |
| TP53 (p53) | 1,00 | 1,22 |
| Autophagy Induction by Intracellular Pathogens: | | |
| EIF1AK3 | 1,00 | 1,32 |
| LAMP1 | 1,00 | 1,14 |
| Autophagy in Response to Other Intracellular Signals: | | |
| CTSD | 1,00 | 1,58 |
| CTSS | 1,00 | 1,91 |
| DRAM2 (TMEM77) | 1,00 | 1,19 |
| EIF4G1 | 1,00 | 1,11 |
| GAA | 1,00 | 1,44 |
| HGS | 1,00 | 0,80 |
| MAPK14 (p38ALPHA) | 1,00 | 1,14 |
| PIK3C3 (Vps34) | 1,00 | 0,93 |
| PIK3R4 | 1,00 | 0,95 |
| RPS6KB1 | 1,00 | 1,08 |
| ULK2 | 1,00 | 1,19 |
| UVRAG | 1,00 | 1,39 |
| Chaperone-Mediated Autophagy: | | |
| HSP90AA1 | 1,00 | 0,55 |
| HSPA8 | 1,00 | 0,66 |
