## Supplementary Figure Legends for "RIT2 reduces LRRK2 kinase activity and protects against alpha-synuclein neuropathology"

**Supplementary Materials**

Supplementary Figures

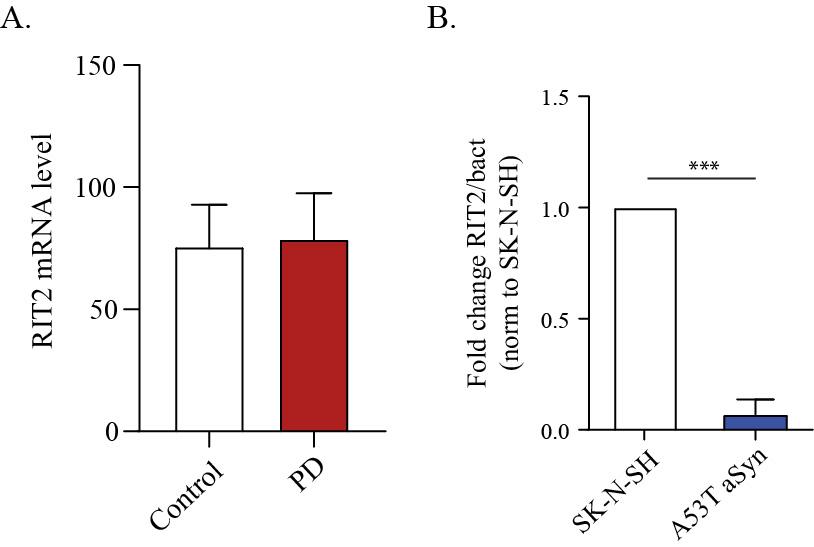

**Figure S1:** *RIT2 gene expression brain tissue and neuroblastoma cell lines.* A) RIT2 mRNA levels are not altered in brain tissue of sporadic PD patients, when compared to controls (GSE7621, controls=9, PD=16). B). RT-qPCR was carried out to assess *RIT2* mRNA levels in recombinant neuroblastoma cell lines. Fold change of RIT2 mRNA is reduced in A53T-aSyn overexpressing cells (n=3). Data are means±SEM of 3 independent experiments. ***p<0.001, two-tailed Student’s t-test.

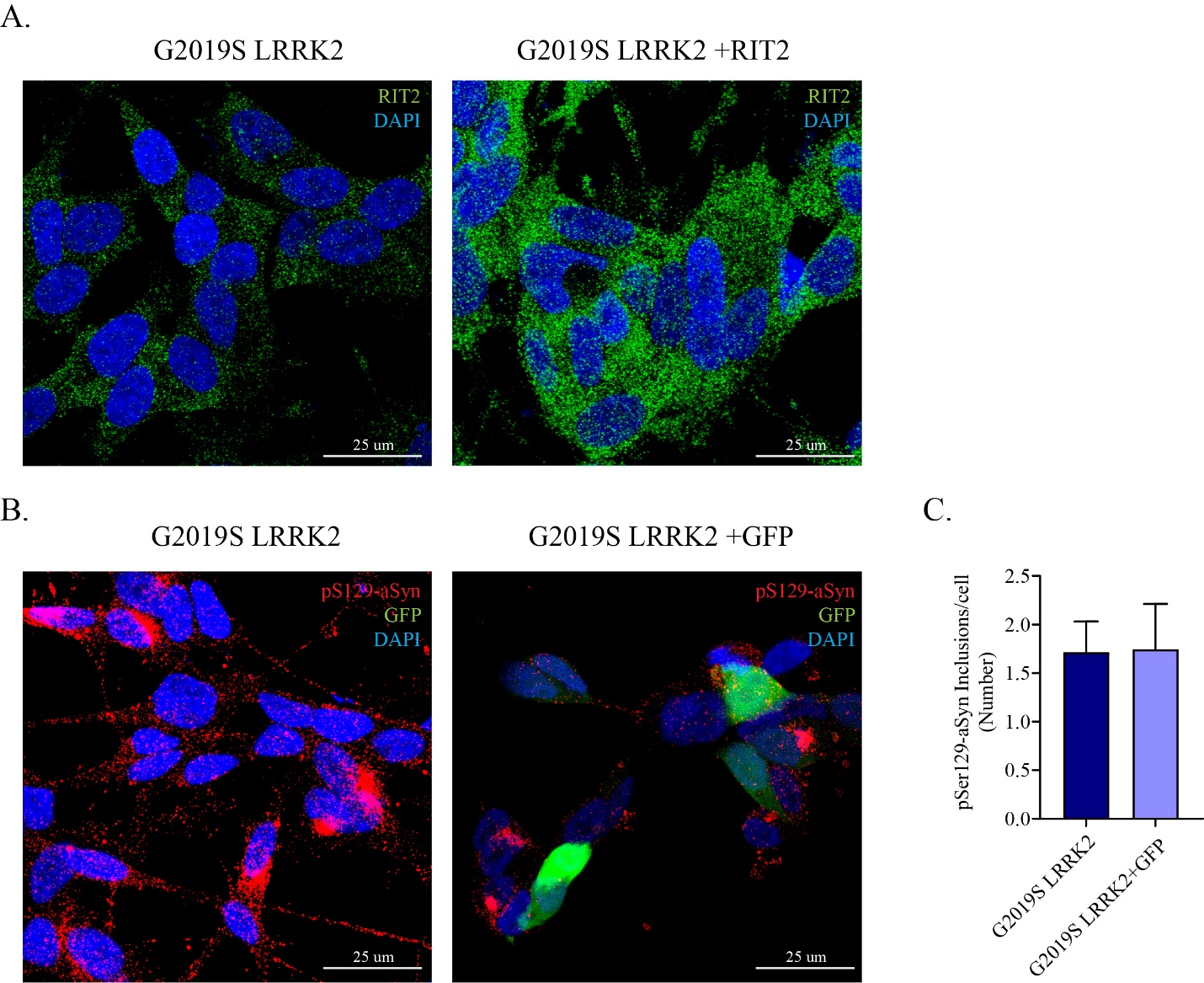

**Figure S2:** *Transfection of RIT2 constructs in G2019S LRRK2 cells.* A) ICC for RIT2 was used to control for nucleofection efficiency. B) Representative images for pS129-aSyn immunostaining and GFP expression in GFP and RIT2-GFP transfected G2019S-LRRK2 cells. C) Quantification of pS129-aSyn inclusions. Expression of RIT2-GFP significantly reduces pS129-aSyn inclusions in G2019S-LRRK2 cells.

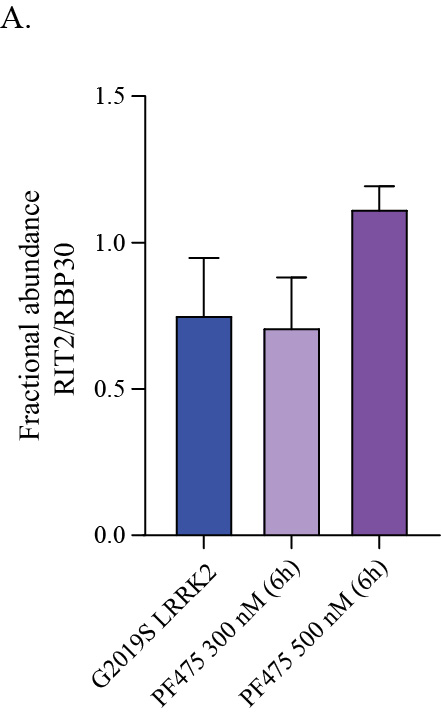

**Figure S3:** *RIT2 gene expression is not changed in PF475-treated G2019S LRRK2 cells.* RIT2 mRNA levels were measured using ddPCR and are presented as fractional abundance of gene of interest over housekeeping gene. (n=3).

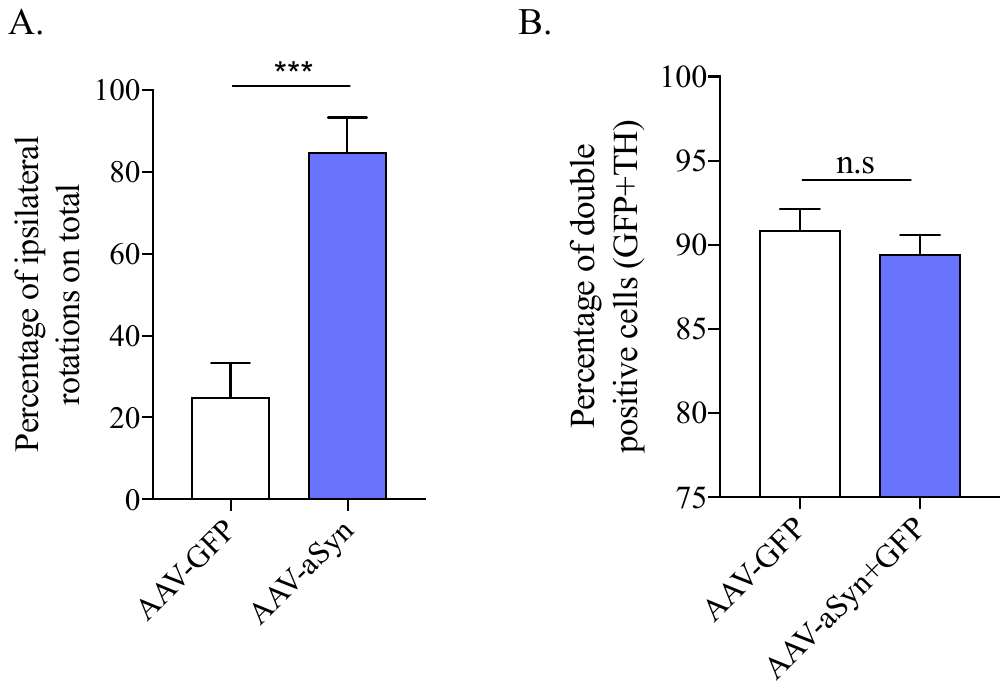
**Figure S4:** *Overexpression of aSyn increases ipsilateral rotations in the amphetamine test and doesn’t affect TH expression level in DA neurons.* A) aSyn overexpression induces a significant increase in ipsilateral rotations in the cylinder with amphetamine test (n^GFP^=7, n^aSyn^=9) ***p<0.001, unpaired two tailed Students’s t-test. B) Overexpression of aSyn in the SNc doesn’t affect TH expression. Around 90% of the cells in the ipsilateral side of AAV-GFP alone or AAV-GFP+aSyn injected mice are double positive for GFP and TH (5 animals/group). Data represented as mean ± s.e.m.

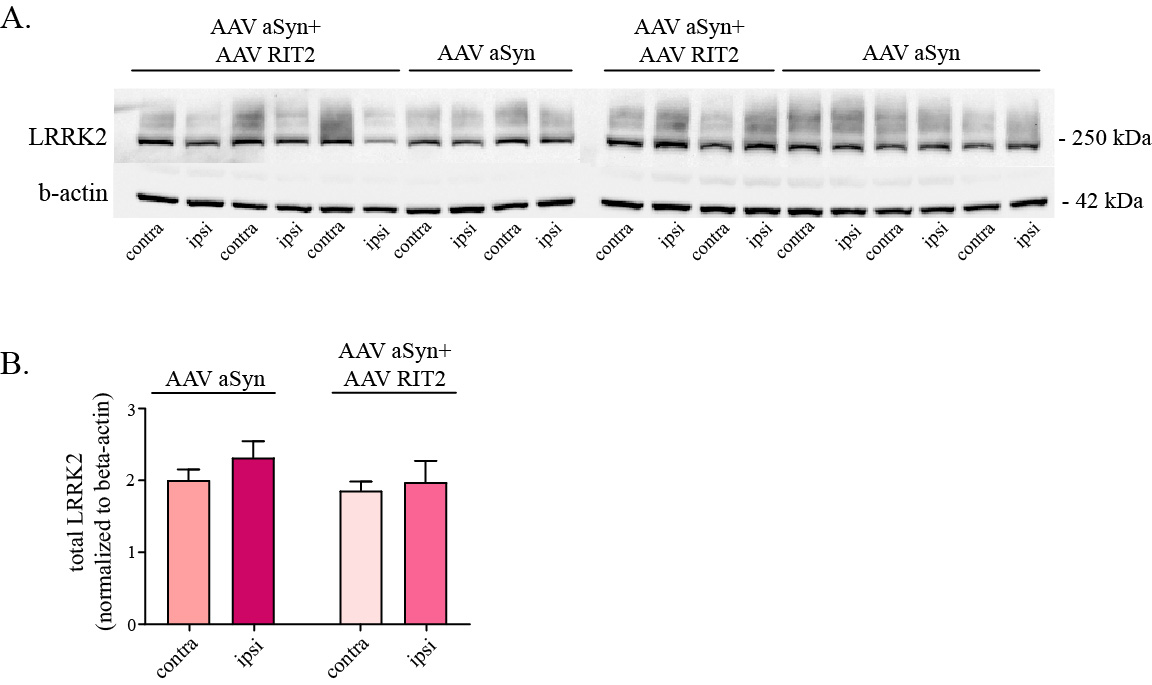

**Figure S5:** *Total LRRK2 levels are not altered by viral overexpression of aSyn and RIT2.* A) Total LRRK2 levels were assessed by Western blot analysis in AAV-GFP, AAV-aSyn and AAV-aSyn+AAV-RIT2 injected mice. B) Total LRRK2 protein levels are not altered by viral overexpression of aSyn and RIT2 (normalized to b-actin) (5 animals/group).
